# Cryo-correlative light and electron tomography of dopaminergic axonal varicosities reveals non-synaptic modulation of cortico-striatal synapses

**DOI:** 10.1101/2025.02.07.637058

**Authors:** Paul Lapios, Robin Anger, Vincent Paget-Blanc, Esther Marza, Vladan Lučić, Rémi Fronzes, Etienne Herzog, David Perrais

## Abstract

Dopamine is an essential brain neuromodulator involved in reward and motor control. Dopaminergic (DA) neurons project to most brain areas, with particularly dense innervation in the striatum. DA varicosities bind to target striatal synapses and form dopamine hub synapses (DHS). However, the basic features of dopamine release sites are still largely unknown. Here we studied the ultrastructure of fluorescent DA and glutamatergic (GLU) synaptosomes isolated from the striatum of adult mice with cryo-correlative light and electron microscopy and cryo-electron tomography. We observed that DA synaptosomes display ∼10 times fewer vesicles than GLU ones. DA vesicles are bigger and less round. Vesicle organization at single nanometer scale indicates that most GLU synaptosomes have tethered and primed vesicles, indicative of a readily releasable pool, while only 39 % of DA synaptosomes have tethered vesicles, which appear not to be primed. In addition, GLU terminals contacted by DA terminals in DHS have more primed vesicles than others. While DA varicosities do not form genuine synapses, their adhesion to cortico-striatal synapses may convey a local regulation of synaptic release properties.

## Introduction

Neuromodulation adjusts network activity and has major impacts on behavior. Among neuromodulators, dopamine acts in the basal ganglia network to encode reward prediction and participate to the initiation of movement. Dopaminergic (DA) projections to basal ganglia originate from two midbrain nuclei, the substantia nigra and the ventral tegmental area. These projections densely innervate the striatum to regulate the activity of spiny projections neurons (SPNs), which are central to functions like motor control, reward prediction and motivation ^1^. Yet, at the ultrastructural level, the organisation of DA transmission is not clear. This situation is in stark contrast with the detailed characterization of neurotransmission machineries at forebrain glutamatergic (GLU) synapses.

At GLU terminals, synaptic vesicles (SVs) are organized in a cluster polarized towards a portion of the plasma membrane called the active zone (AZ) which faces the post-synaptic density (PSD). The AZ contains specific proteins such as RIM1/2, bassoon and ELKS ^2^. Prior to fusion, proximal SVs follow a series of steps, which can be observed with cryo-electron tomography (cryo-ET), from initial tethering, where SVs are tethered to the plasma membrane by one filament of approximately 10-25 nm ^3,4^, to the formation of multiple short tethers, likely comprising the SNARE complex, which brings SVs closer than 5 nm to the plasma membrane ^5^. These steps strongly depend on the AZ proteins RIM1 and Munc13a, which are known to control fast SV exocytosis ^6^. Thus, the presence of docked and primed vesicles may constitute a hallmark of readily releasable vesicles in axons.

Dopamine is also exocytosed from vesicles within milliseconds after stimulation ^7,8^. This process relies on the calcium sensor Synaptotagmin-1 ^9,10^. Dopamine release shows strong paired pulse depression which is in part controlled by Synaptotagmin 7 ^11^. Moreover, DA axons contain AZ proteins, RIM1, Munc13 and ELKS, which are important for fast dopamine release ^12,13^. However, only ∼30 % of DA varicosities contain such assemblies ^12^. This observation is supported by functional evidence showing that around a quarter of dopaminergic varicosities are active, as assessed by release of fluorescent neurotransmitter analogues ^14^. Furthermore, serial electron microscopy studies of dopaminergic axon terminals revealed heterogeneous vesicular content, with terminals containing varying combinations of small and large vesicles, while some terminals appeared to lack vesicles altogether ^15^. Overall, DA terminals exhibit a non-stereotyped vesicular organization which may explain the functional diversity of release. However, observations of the accurate spatial arrangement of DA vesicles prior to fusion has been hampered by chemical fixation, staining procedures and difficulties to identify rare DA axons which negatively affects the preservation of cellular morphology and precludes interpretation of molecular details ^16^.

The relationship between DA release sites and target cells remains poorly documented. Upon release, dopamine binds to G-protein coupled receptors of the D1 group (D1/5R) or D2 group (D2-4R) whose signals respectively increase or decrease the excitability of target cells. On pre-synaptic terminals they influence SV release probability, while at the post-synapse they act on ion channels and glutamate receptors (reviewed in ^17^). In particular, neurons projecting from motor and prefrontal cortical regions form GLU cortico-striatal (CS) synapses responsible for the activation of SPNs. Interestingly, the stimulation of dopamine release in acute slices attenuates the release kinetic from a subset but not all GLU terminals ^18^, showing that dopamine influences the activity of CS synapses. However, the precise ultrastructure through which dopaminergic terminals interact with their synaptic targets is unclear. DA synapses showing classical pre and post-synaptic features represent a minority ^15,19–21^. Nevertheless, DA boutons are often found in close apposition with either pre or post-synaptic elements of CS synapses ^22^. This proximity is functionally relevant, because DA release generates 2 µm wide hotspots of dopamine as estimated using carbon nanotube sensors ^23^. Moreover, the synchronisation of DA phasic release with local glutamate uncaging induces structural plasticity at spines ^24^.

Isolation of DA synaptosomes from striatal tissue and sorting by fluorescence activated synaptosome sorting (FASS) revealed that most DA terminals are in close contact with other terminals, such as GLU presynapses ^25^. These interactions are conserved even though tissue homogenisation and droplet-based fluorescence activated sorting exposed structures to significant mechanical shearing forces. We termed these multipartite DA containing synapses “dopamine hub synapses” (DHS). Among them, 25 % were formed with CS synapses marked by the vesicular glutamate transporter VGLUT1 ^22,25,26^. Importantly, CS-DHS contain an increased signal for the presynaptic proteins VGLUT1 and bassoon compared to other CS synapses. Collectively, these findings highlight the importance of a local dopaminergic signalling for the regulation of CS synapses. However, there is still a major lack of ultrastructural observations in close-to-native conditions of DA terminals associated with their target synapses.

Cryo-correlative light and electron microscopy (cryo-CLEM) and cryo-electron tomography (cryo-ET) enable identification of vitrified fluorescently labeled terminals and a close-to-native 3D observations of cellular ultrastructure and protein complexes at a nanometer scale ^27–29^. This allows determination of the vesicular organisation in different synaptic types and characterization of protein complexes ^30^. Synaptosomes are a suitable model for cryo-EM because they can be vitrified by plunge freezing, imaged in transmission EM without thinning, and because they preserve association between terminals and important pre- and postsynaptic function ^16,29,31^. Here, we applied cryo-CLEM and cryo-ET to fluorescently labeled synaptosomes in order to determine the ultrastructural features of DA terminals, CS synapses and CS-DHS. We reveal the spatial organization of DA vesicles, as well as the structural association with GLU synapses in DHS. Importantly, we observed tethered SVs in DA terminals and quantified differences in SV organization of CS that are correlated to the association with DHS.

## Results

### Identification and observation of GLU and DA synaptosomes by cryo-CLEM and cryo-ET

We assessed the ultrastructure of identified glutamatergic (GLU) and dopaminergic (DA) terminals extracted from the striatum of adult mice using cryo-CLEM combined with cryo-ET as described in Figure 1. We prepared synaptosomes from the striata of 12 to 18 week-old mice as previously described ^25^ and detailed in Methods and Figure 1A,B. We used 3 mouse lines. For GLU elements we took advantage of the knock-in (KI) mouse line in which the VGLUT1 open reading frame is tagged with the sequence of the fluorescent protein Venus ^32^. In the striatum, VGLUT1^venus^ specifically labels cortico-striatal terminals ^22,26^. For DA synaptosomes we injected an adeno associated viral vector carrying sequences for Cre-dependent mNeonGreen expression (AAV1 pCAG-Flex-mNeonGreen) in the midbrain of dopamine transporter promoter (DAT)-Cre transgenic mice, which specifically labels dopaminergic neurons ^25^. To identify CS-DHS we used dual tagging of GLU (green) and DA (red) synaptosomes by crossing VGLUT1^venus^ mice with DAT-Cre ^33^ and the reporter line Ai14-tdTomato ^34^. Striatal synaptosomes were incubated at 37 °C for 15 minutes before use. They are capable of depolarization-evoked exocytosis and recycling, as shown by uptake and release of the membrane dye FM4-64 (Figure S1), in accordance with previous work ^35,36^. This suspension was mixed with electron dense fluorescent fiducial beads for alignment and gold beads for tomogram reconstruction. The mixture was applied on an EM grid, plunge frozen into liquid ethane (Figure 1C) and kept in liquid nitrogen for further use.

**Figure 1:**
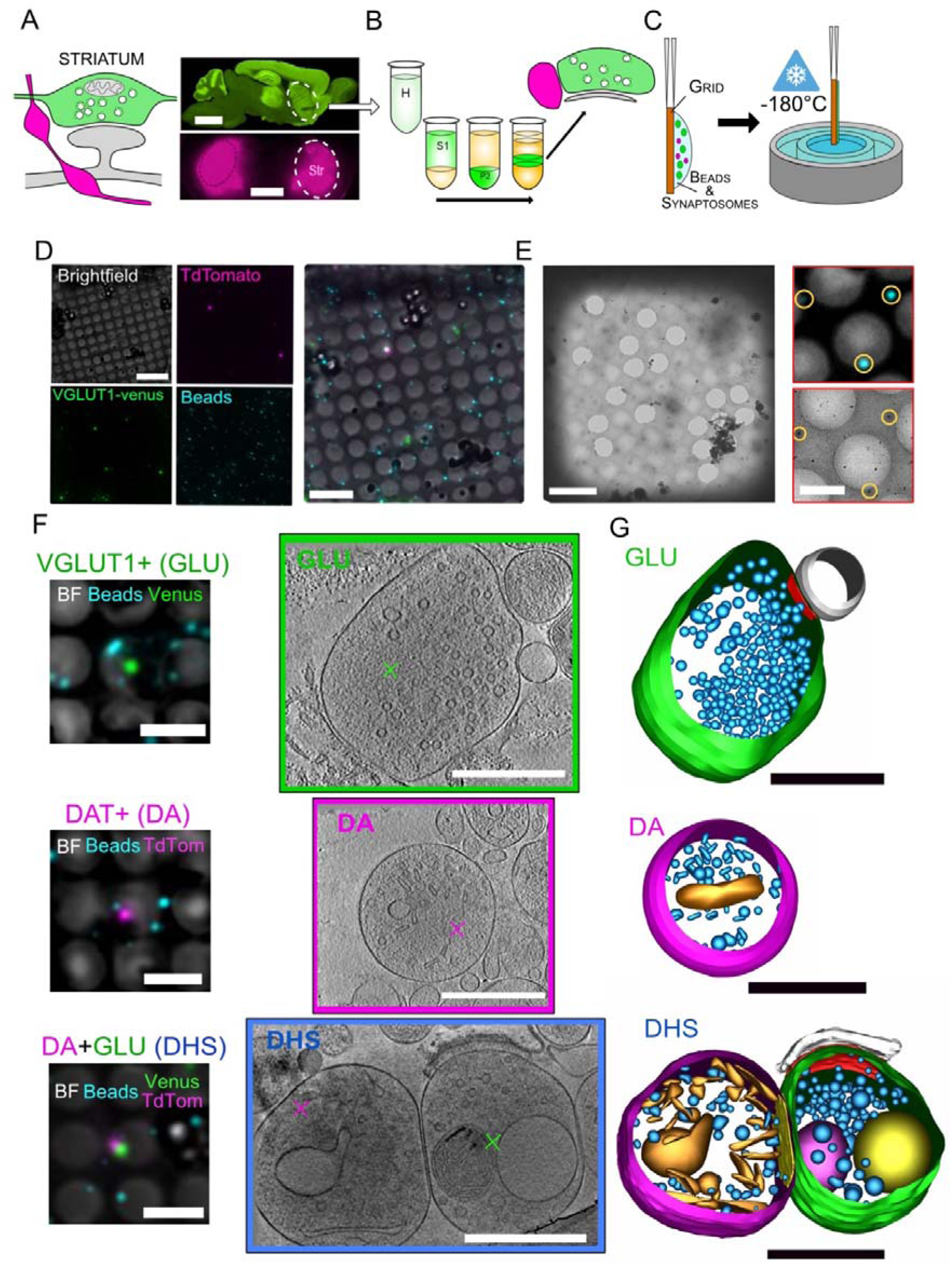
Cryo-CLEM and cryo-ET of cortico-striatal and dopaminergic synaptosomes. **A**, *Left*, schematic of the expected organization between DA varicosities (magenta) and GLU synapses (green), forming DHSs. *Right top*, VGLUT1^venus^ fluorescence in a sagittal slice. CS synapses generate an intense fluorescence signal in the striatum (dotted line). *Right bottom*, DA projections produce a bright striatal fluorescence in a coronal slice of a DAT-Cre mouse injected with the AAV-Flex-mNeonGreen (magenta). Scale bars: 3 mm **B**, Scheme of synaptosomes preparation from fresh striatum obtained after differential centrifugations. The fractions (H, S1, P2, B, see Methods) are increasingly enriched in synaptosomes. After discontinuous density gradient centrifugation, synaptosomes are diluted and centrifuged to remove Ficoll. **C**, A suspension of synaptosomes mixed with fluorescent and gold fiducial beads is layered on the grid, blotted and plunge-frozen in liquid ethane. **D**, Example of a cryo-fluorescence microscopy image of a single grid square from a VGLUT1 ^venus^ x DAT-Cre x Ai14-tdTomato with 4 channels: bright field, fluorescence of tdTomato, Venus and blue fiducial beads. These images were used to select synaptosomes of interest, green (VGLUT1^venus^) or magenta (DAT+) next to several blue beads, for observation with electron microscopy. Scale bar 10 µm. **E**, Left cryo-EM image of the same square as displayed in D. Scale bar 10 µm. Right, subset of cryo-fluorescence (top) and EM (bottom) of the same portion of grid with 3 fiducial blue beads visible in both modalities (yellow circles) for fine registration. Scale bar: 3 µm. Synaptosomes with clearly visible beads and good ice quality were further imaged with high magnification tilted series. **F**, Reconstructed tomograms of fluorescent synaptosomes obtained by cryo-electron tomography. *Left*, overlay of fluorescence and bright field channels showing tagged synaptosomes of interest and fiducial beads. Scale bars: 5 µm. *Right*, representative single tomographic slice of 1.558 nm of thickness showing a clear structure outline and intracellular organelles. Magenta or green cross marks point to the registered centre of the fluorescent tag (GLU or DA synaptosome), Scale bars: 500 nm. **G**, 3D models from the synaptosomes displayed in F obtained by segmentation of membranes. The display of 3D models are as follows: plasma membrane of synaptosomes in green for GLU or magenta for DA; Small (synaptic) vesicles in blue; elongated endoplasmic-reticulum-like structures in orange; large endosome-like organelles in yellow; mitochondrion in magenta. For both GLU synaptosomes, a clear synaptic cleft was detected, with either a closed post-synaptic element (PSE) in grey (top), or an opened post-synaptic membrane (bottom). Both of them exhibit a post-synaptic density. The presynaptic GLU active zones are shown in red. Scale bar: 500 nm

We first observed the grids with a cryo-fluorescence microscope. We selected isolated fluorescent spots with nearby fiducial beads for alignment (Figure 1D). In mice with both fluorescent markers for GLU and DA synaptosomes, we selected either an individual isolated spot, or pairs of spots where the two colours were separated by less than 1 µm corresponding to putative DHS. We then observed the same grids with a cryo-transmission electron microscope (Talos Arctica), and located regions of interest based on bead patterns (Figure 1E). We used the images of neighbouring independent fiducial beads to measure the pointing precision between cryo-fluorescence and cryo-electron microscopy (Figure S2; 117 ± 82 nm, n = 64). This pointing precision is smaller than the radii of GLU and DA synaptosomes (see below). Therefore, we can identify unambiguously the terminals of interest in cryo-EM, either GLU or DA. Finally, we acquired tilt series of selected synaptosomes and reconstructed tomograms with weighted back projection method using IMOD ^37^. We obtained tomograms from 19 preparations (Table1): 4 from VGLUT1-Venus mice (39 GLU synaptosomes), 5 from DAT-Cre+AAV-mNeonGreen mice (43 DA synaptosomes) and 10 from VGLUT1-Venus*DAT-Cre-Ai14-tdTomato (62 GLU and 51 DA synaptosomes, among which 32 form DHSs). We thus have a dataset of tomograms of 101 GLU synaptosomes and 94 DA synaptosomes with 32 CS-DHSs (Figure 1F). We created 3D models of these synaptosomes, in which we segmented the plasma membrane of the synaptosome, internal membranes and, when applicable, identified adhering structures such as a post-synaptic element (PSE), as illustrated in Figure 1G.

### GLU synaptosomes are larger and contain more SVs than DA synaptosomes

The mean size of GLU synaptosomes, as measured by their maximal extension, is 823 nm, which corresponds to a visible volume of 0.0915 µm^3^ (Figure 2A,C,D). They all contain small round SVs (Figure 2A). Each synaptosome contains 13 to 872 vesicles, 191 on average (Figure 2E), for an average density of 1956 vesicles/µm^3^ (Figure 2F). In addition, 21/101 synaptosomes contained a mitochondrion (Figure 1F, 2A) and some contained large organelles or pleiotropic organelles such as round vacuoles, multivesicular bodies, as well as other intracellular features, such as filaments and clathrin-coated vesicles (Figure S3A-F). The majority of GLU presynaptic elements (64/101, 63%) are connected to a PSE, which also contains occasionally intracellular organelles (Figure S3G-I). A gallery of representative tomographic slices and 3D models of GLU synaptosomes is shown in Figure S4.

**Figure 2:**
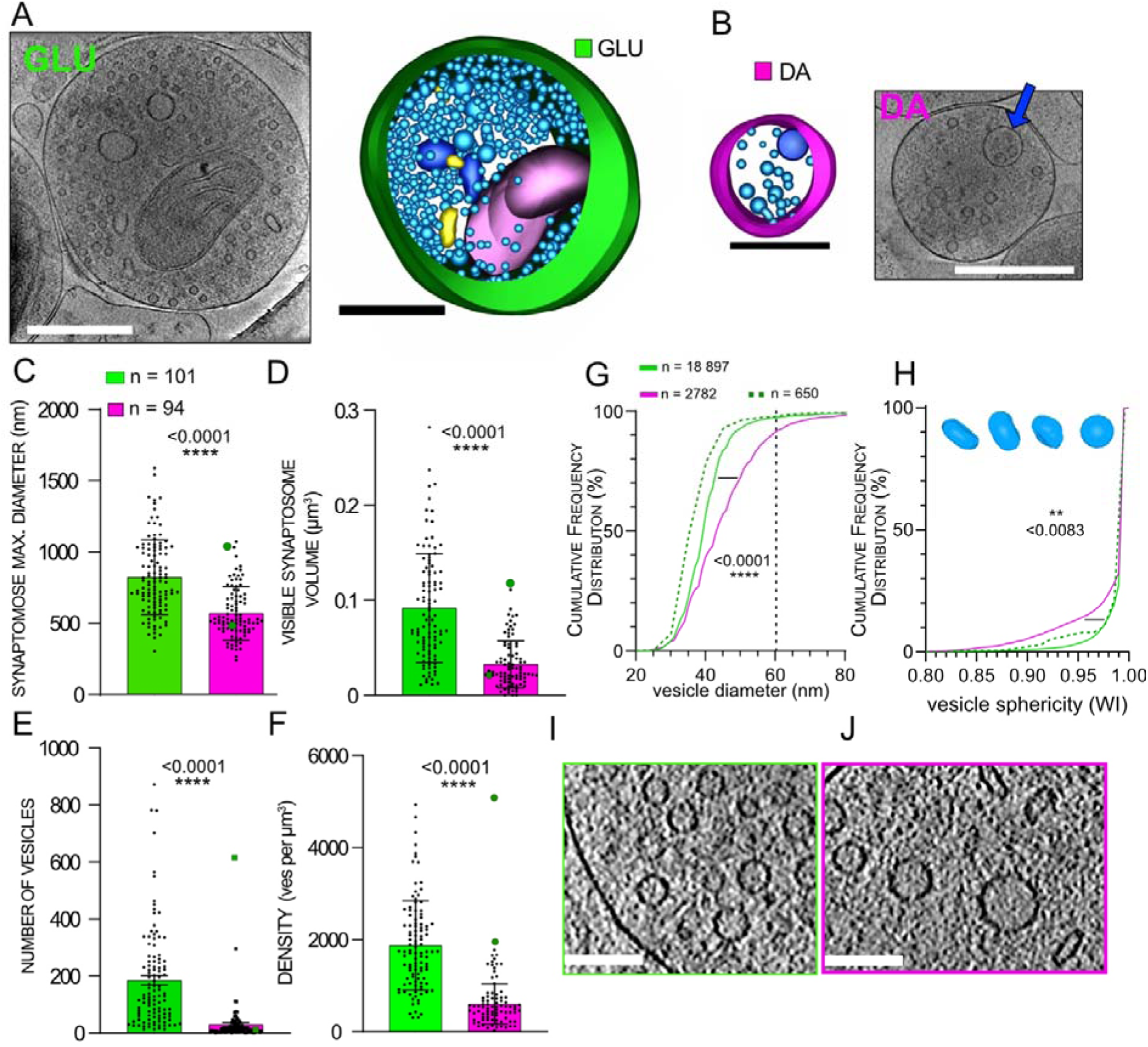
Comparison of synaptosome size and vesicular content for GLU and DA synaptosomes. **A,B**. Example of a single tomogram plane of 1.558 nm thickness, and corresponding to 3D models of GLU (A) and DA (B) synaptosomes. Same color code as in Figure 1. In addition, multivesicular bodies are visible in the tomogram slices (blue arrows) and in the models (dark blue). Scale bars 500 nm **C-F**, Distributions and average diameters (C), visible volumes (D) number of vesicles (E) and vesicle density (F) of 101 GLU and 94 DA synaptosomes. The distributions are all different (t test p < 0.0001). **G**, Cumulative distribution of vesicle diameters in GLU (green line, n = 18897) and DA (magenta line, n = 2782) synaptosomes. The diameters are significantly different (Kolmogorov-Smirnov test p < 0.0001). The green dotted line (n = 650) corresponds to vesicles in DA terminals with a PSE sharing the features of asymmetric synapses (examples in S6). **H**, Cumulative distribution of vesicle sphericity (WI). GLU vesicles are significantly more spheric than DA vesicles (Kolmogorov-Smirnov test p < 0.0083). We show examples of DA vesicles with WI = 0.85, 0.89, 0.95 and 0.99 from left to right above the plot. **I**,**J**, Cropped zoomed in tomograms of GLU (I) and DA (J) synaptosomes display the variety of vesicle sizes and shapes. Scale bars 100 nm.

DA synaptosomes (example Figure 2B) have a mean size of 575 nm, which corresponds to a visible volume of 0.0341 µm^3^, significantly smaller than GLU synaptosomes (Figure 2C,D, p-value < 0.0001). DA synaptosomes contain on average 30 vesicles but their number is quite variable: some are almost empty (23/94 synaptosomes have less than 5 vesicles) while others contain tens or even hundreds of vesicles (Figure 2E and Figure S5, S6A-B). Overall, the vesicle density of DA synaptosomes is about 3-fold smaller (723 per µm^3^) than for GLU synaptosomes (Figure 2F). DA synaptosomes also contain other organelles, such as vacuoles, mitochondria, multivesicular bodies (Figure 1F, 2B, S6C-H). Table 2 reports the number of DA synaptosomes with these organelles. While SVs in GLU synaptosomes are spherical, small and uniform in size (outer diameter 39.8 ± 5.8 nm, n = 18 897, with only 2.7% vesicle larger than 60 nm), vesicles in DA synaptosomes are significantly larger (44.9 ± 9.9 nm, n = 2782, p < 0.0001) with 8.8% of vesicles having a diameter in the 60-100 nm range (Figure 2G). In total, 66/94 DA synaptosomes have at least one vesicle bigger than 60 nm, with a proportion of 23.5 ± 19.6 % big vesicles in these synaptosomes. Moreover, vesicles in DA synaptosomes are often elongated or pleomorphic (Figure 2H,J). We used Wadell’s index WI (see Methods) to quantify vesicle sphericity. Most vesicles in GLU terminals have an index close to 1 (perfect sphere). However, vesicles in DA synaptosomes are significantly less round than the ones in GLU synaptosomes (Figure 2H, p-value = 0.0083). These elongated vesicles are spread across the synaptosomes (Figure 2J, S5). Overall, there are 65/94 DA synaptosomes containing at least one elongated vesicle (WI < 0.95) (Figure 2J). In these synaptosomes there are 29.2 ± 26.9 % of elongated vesicles.

### Contact zones between GLU, DA synaptosomes and PSEs

In GLU synaptosomes we identified a clear PSE separated from a presynaptic terminal by a clearly defined synaptic cleft in 64/101 (63%) synaptosomes. Typically, the PSE is defined by a sealed plasma membrane compartment (Figure 1F,3A,C, S4) but in some cases it had an open membrane adhering to the presynaptic terminal forming a synaptic cleft (Figure 1F,S4). The area of contact between GLU and PSE is characterized by roughly parallel membranes defining a wide cleft (31.8 ± 6.2 nm, n = 34) with dark material around the midline (Figure 3D,E), as observed previously ^38,39^. The active zone (AZ), defined as the membrane region of GLU synaptosome in contact with the PSE (outlined in red in Figure 3A), has an area of 0.079 ± 0.055 µm^2^ (n = 33) (Figure 3F). We analysed the presence of protein density in the pre- and post-synaptic elements of GLU synaptosomes. The normalized densities decrease by 20 % in both compartments (Figure 3J,K). However, we detected a clear increase in density 10 to 25 nm from the PSE membrane, which corresponds to the post-synaptic density (PSD), as detected with other modalities of electron microscopy for synapses in situ ^40^ and synaptosomes with cryo-ET ^29^. In contrast, only two out of 94 DA terminals were separated by a wide cleft from a characteristic PSE containing PSD, as in GLU synaptosomes. Interestingly, one of these DA synaptosomes has the most vesicles (610), all of them small and round, which makes it indistinguishable from classical GLU synaptosomes (Figure S6A). The other one has 40 vesicles (Figure S6B). In these two synaptosomes, the distribution of vesicle sizes and sphericity matches distributions seen for GLU synaptosomes (dotted line in Figure 2G,H). Another DA synaptosome has 295 vesicles (see two upper dots in Figure 2E) and also resembles classical GLU synaptosomes, even though it was not connected to a PSE but engaged into a CS-DHS. These DA synaptosomes could originate from neurons co-expressing VGLUT2 and which are able to release glutamate and generate AMPAR-mediated post-synaptic potentials ^41,42^.

**Figure 3:**
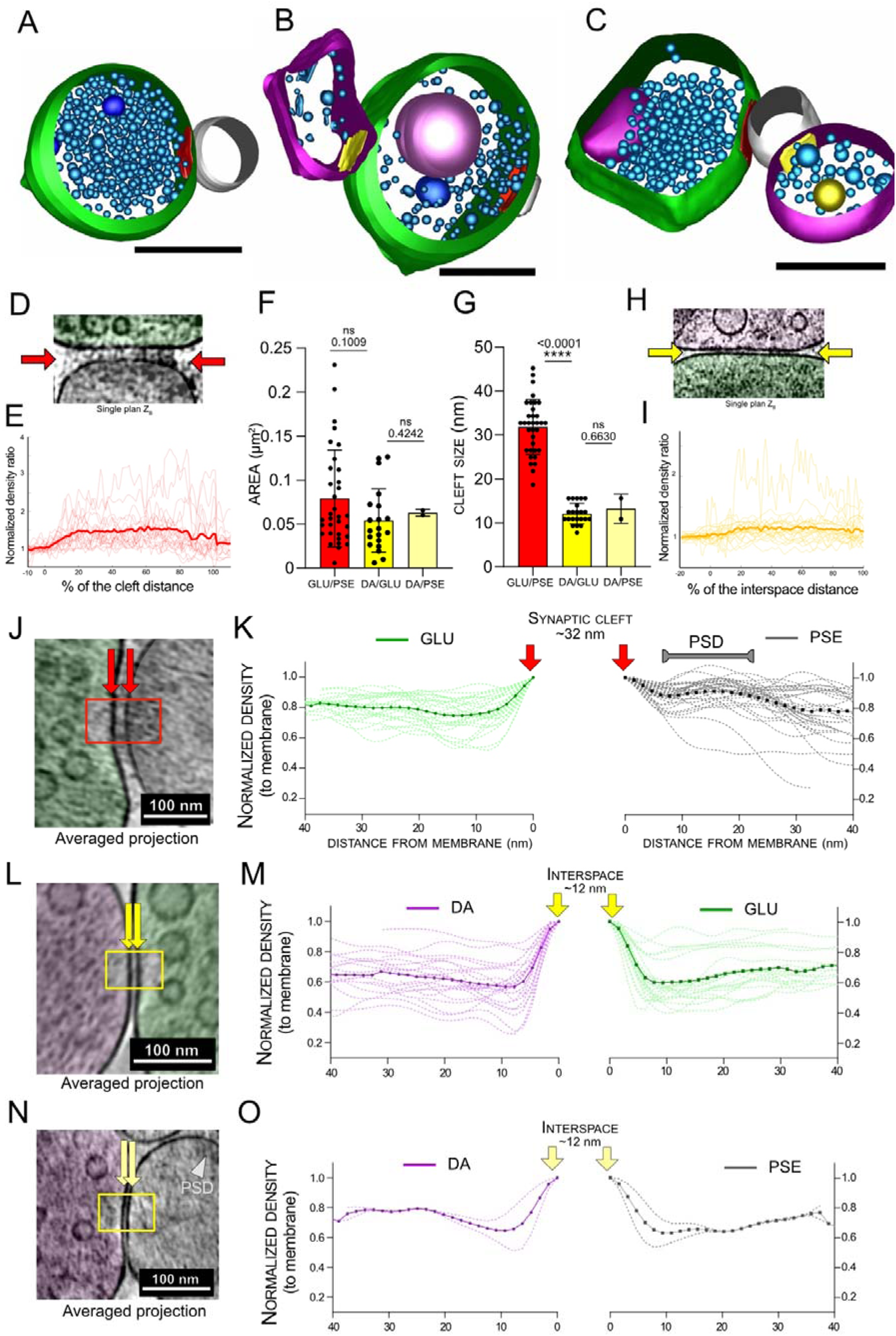
Contact zones between GLU, DA synaptosomes and PSEs. **A-C**, 3D models of a GLU presynaptic elements (green) connected to a PSE (grey). The active zones are drawn in red. In B, a DA element connects the GLU presynaptic element. In C, a DA element connects the PSE. The contact zones with DA are drawn in yellow. **D**, single plane showing the synaptic cleft between a GLU synaptosome (outlined in green) and a PSE (outlined in grey). The electron dense material in the middle of the cleft is clearly visible (red arrows). **E**, Pixel intensities along the length of the synaptic cleft, normalized to area outside the synaptic cleft (n = 20). **F**, Area of active zones of GLU synaptosomes and contact zone between DA and GLU synaptosomes or DA and PSEs. **G**, Membrane to membrane distance of the cleft of: GLU with PSE (synaptic cleft), DA with GLU synaptosomes, and DA with PSE. **H**, Single plane showing the cleft between a DA (magenta) and a GLU synaptosome (green). **I**, Pixel intensities along the length of the DA/GLU contact site, normalized to region outside the contact site (n = 24). **J**, Image of a contact site (synaptic cleft) between GLU and PSE. **K**, Normalized intensity profiles in the cytoplasm starting 5 nm away from the plasma membrane, towards the center of GLU (green) and PSE (grey). In both cases, the averaged intensity reached a plateau around 80% 40 nm away. In the PSE, the intensity clearly increases 20 nm away, which corresponds to the PSD. We observe no clear increase on the presynaptic side of GLU. **L-M**, Same as J-K for DA-GLU contact sites. **N-O**, Same as J-K for DA-PSE contact sites.

Nevertheless, many DA terminals are in close apposition with other structures which are occasionally resembling a GLU terminal (Figure S5A) or other, harder to characterize structures which could be parts of postsynaptic spines. To get a better insight on the features of contact zones between DA and GLU terminals, we analyzed 32 tomograms from apposed GLU and DA synaptosomes which correspond to CS-DHSs ^25^. The area of contact between GLU and DA synaptosomes have a similar size as GLU active zones (Figure 3B,F). The width of the cleft between GLU and DA terminals is 12.1 ± 2.4 nm, much smaller than the synaptic cleft of GLU synaptosomes (Figure 3G, p < 0.0001). Along this cleft, dense material is also observed, albeit not as pronounced as for GLU/PSE synaptic clefts (Figure 3H,I). The intracellular side of DA and GLU terminals around the adhesion site are devoid of visible densities such as PSDs (Figure 3L,M). In 2 tomograms, the DA terminal is not in contact with the presynaptic GLU synaptosome but with the PSE (Figure 3C,S7B). The contact area (0.062 and 0.066 µm^2^) and cleft sizes (10.9 and 15.6 nm) are in the same range as for DA/preGLU contact sites (Figure 3F,G). Similarly, the densities around the contact site are in the same range, without signs of a PSD (Figure 3N,O). More examples of DHS virtual planes and models are available in Figure S7.

### Vesicles in GLU and DA terminals are tethered to the plasma membrane and are interconnected

Synaptic vesicles in GLU and GABAergic synapses are divided into functional pools ^43^. SVs which fuse first with the plasma membrane upon calcium stimulation comprise the readily releasable pool, which was defined morphologically by EM of chemically fixed, dehydrated samples as the vesicles which appear docked, that is in direct contact with the plasma membrane ^44,45^. Cryo-ET revealed that in close to native state, SVs are not docked but are tethered to the plasma membrane via one or several electron dense filaments, as observed in synaptosomes and dissociated neuronal cultures ^3,4,29,30,46^. We found at least one tethered vesicle at the active zone in almost all quantified striatal GLU synaptosomes (24/26, 92 %) (Figure 4A). Tethers, as well as connectors (bridges that interconnect SVs) were detected using an automated, template-free method by the hierarchical connectivity algorithm ^47^. Among proximal vesicles, defined as those localized less than 45 nm from the plasma membrane, 40 ± 7 % have one or more tethers (Figure 4C). These tethers have a length of 13.9 ± 7.5 nm (Figure 4D). The number of tethers increases as vesicles are located closer to the plasma membrane (Figure 4E). Remarkably, vesicles located less than 5 nm from the plasma membrane have on average 3 tethers, which corresponds to synaptic vesicles that are functionally primed for fusion^3,4^.

**Figure 4:**
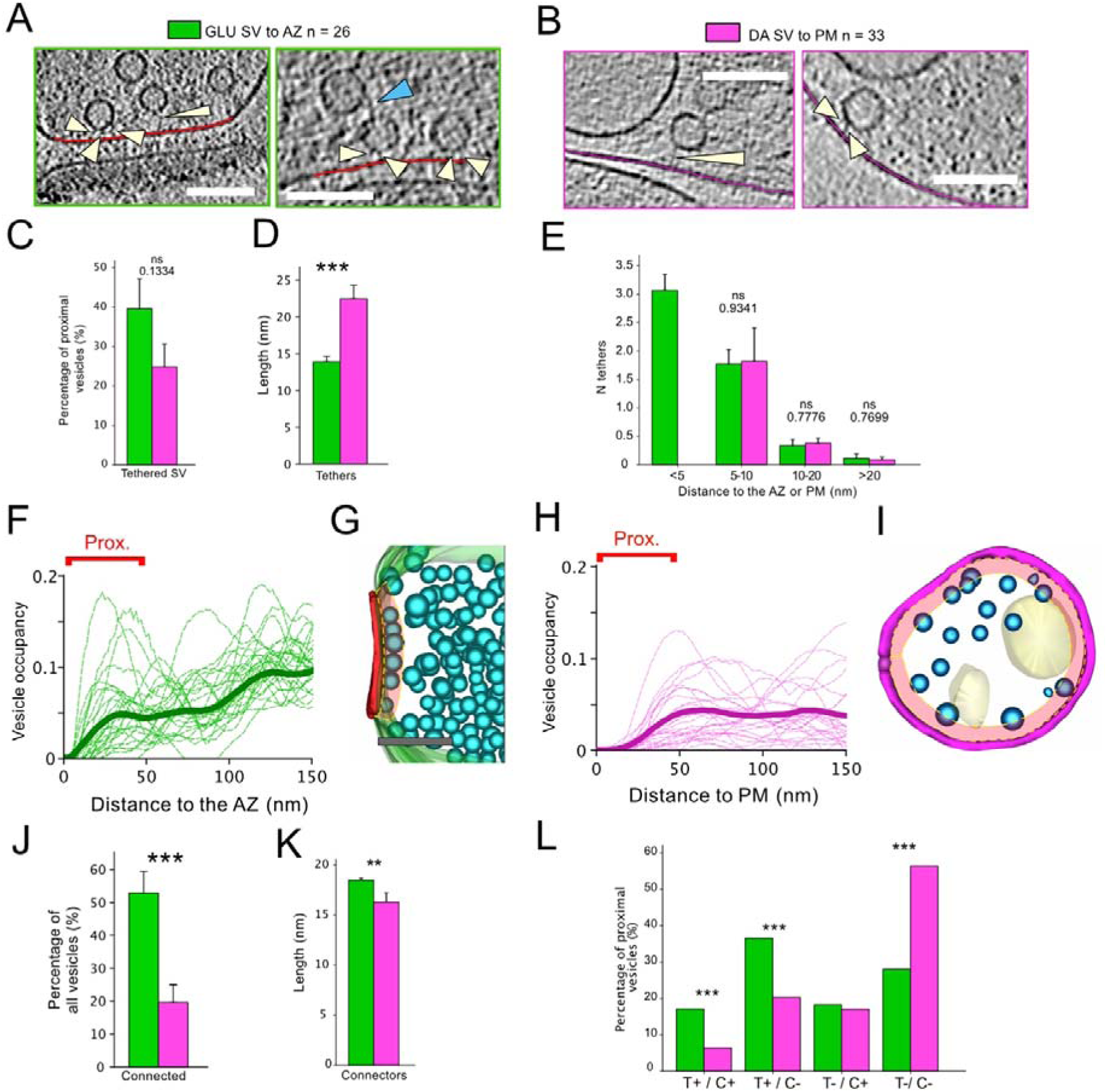
Vesicles with tethers and connectors in GLU and DA synaptosomes. **A**, Tomogram planes showing vesicles tethered to the plasma membrane in GLU synaptosomes (white arrowheads). Some vesicles are connected via multiple tethers. Connectors between vesicles are also visible (light blue arrowheads). Scale bar 100 nm **B**, Same as A for DA synaptosomes. **C**, Percentage of proximal vesicles (located less than 45 nm) connected by at least one tether in GLU and DA synapses (t-test; p-value = 0,133). **D**, Length of tethers connecting vesicles to the plasma membrane in GLU and DA synapses (t-test; p-value < 0.001). **E**, Average number of tethers for vesicles at various distances from the plasma membrane (t-tests; p-values = 0.934; 0.777; 0.770). **F**, Fraction of volume occupied by vesicles vs distance to the AZ for GLU synapses. The mean distribution is shown as a thick line. **G**, 3D model of a GLU synaptosome around the active zone showing the concentration of proximal vesicles that are tethered. **H**, Fraction of volume occupied by vesicles vs distance to the plasma membrane for DA synapses. The mean distribution is shown as a thick line. **I**, 3D model of a DA synaptosome with the zone of proximal vesicles. **J**, Percentage of all vesicles connected to another in GLU and DA synaptosomes (t-test; p-value = 0.0004). **K**, Average length of connectors (t-test; p-value = 0.0021). **L**, Proportion of proximal vesicles that are tethered and/or connected in GLU and DA synaptosomes (t-test; p-value = 0.0008).

In DA synaptosomes, we found clear examples of tethers between vesicles and the plasma membrane, as well as vesicles connected by more than one tether (Figure 4B). We detected vesicles tethered to the plasma membrane in 39 % (13/33) of DA synaptosomes. The percentage of tethered vesicles among proximal vesicles in DA is 25 ± 6 %, which was lower, but not significantly different from GLU synaptosomes (Figure 4C). However, tether length is on average significantly larger in DA versus GLU (DA: 22.43 nm; std. 12.21; GLU: 13.89 nm; std: 7.47; p-value < 0.001) (Figure 4D) likely a direct consequence of the complete absence of tethered DA vesicles localized less than 5 nm from the plasma membrane (Figure 4E). Importantly, for DA and GLU vesicles located further away, the number of tethers per SV is not different (Figure 4E).

Another morphological hallmark of the polarized exocytosis of SV in cortical synaptosomes is the larger fraction of volume occupied by proximal synaptic vesicles towards the active zone ^3,4,29^. GLU synaptosomes showed a mean peak occupancy around 20 nm from the plasma membrane that is clearly visible in individual occupancy profiles (Figure 4F,G) and is similar to those observed for cortical synaptosomes ^3,4,29^. In DA synaptosomes, the occupancy lacked the peak in the proximal vesicles region, but occasional enrichment close to the whole plasma membrane was observed (Figure 4H,I). However, the absence of a morphologically identifiable DA active zone, may have biased the measurement of occupancy, because it included a larger, possibly irrelevant volume in our analysis.

The percentage of inter-connected SVs was significantly higher in GLU (53%) than in DA synaptosomes (20%) (Figure 4J) and the length of connectors was significantly different (DA: 18.50 nm; std. 9.18 and GLU: 16.33 nm; std. 13.07; p-value = 0.0021) (Figure 4K). Finally, among proximal vesicles, there were more SVs that were both tethered and connected in GLU (17%) than in DA synaptosomes (6.4%). In the latter, the majority of vesicles were neither tethered nor connected (Figure 4L).

### Tethered SVs highlight a subpopulation of DA terminals

We compared DA synaptosomes that contain (T+) to ones that do not contain (T-) tethered vesicles (Figure 5A). We excluded from the analysis the 2 DA synaptosomes with a PSE (see Figure S6AB). The T+ DA synaptosomes contain significantly (twice) more SVs (Figure 5B), and are significantly larger than the T-synaptosomes (Figure 5C). On average, T+ synaptosomes contain 2 tethered vesicles, and a maximum of 5 (Figure 5D). In T+ DA terminals we do not observe a clear AZ, which corresponds in GLU synapses to the location facing the synaptic cleft where tethered synaptic vesicles concentrate and undergo exocytosis upon stimulation. We looked for a putative DA active zone in two ways. First, the active zone could locate at or right next to the contact zone with GLU terminals. However, tethered vesicles are almost never found at the contact site. Second, an active zone is expected to concentrate tethered vesicles. Therefore, we analyzed DA synaptosomes with multiple tethered vesicles. Among the 13 T+ DA synaptosomes, 6 had multiple tethered vesicles, for a total of 19 tethered vesicles (Figure 5D,E). The shortest arc distance (i.e. staying on the plasma membrane) between these vesicles is on average 210 nm (Figure 5F). This is smaller than the average distance between random points on the synaptosome plasma membrane (estimated at 567 nm, see Methods for derivation). This suggests that tethered DA vesicles may gather in a preferential zone of the plasma membrane, a putative active zone. Moreover, in T+ DA synaptosomes, vesicles are located closer to the plasma membrane than in T-DA synaptosomes (Figure 5G), reinforcing the idea that in DA synaptosomes vesicle location is polarized to a putative active zone where vesicle may tether and fuse.

**Figure 5:**
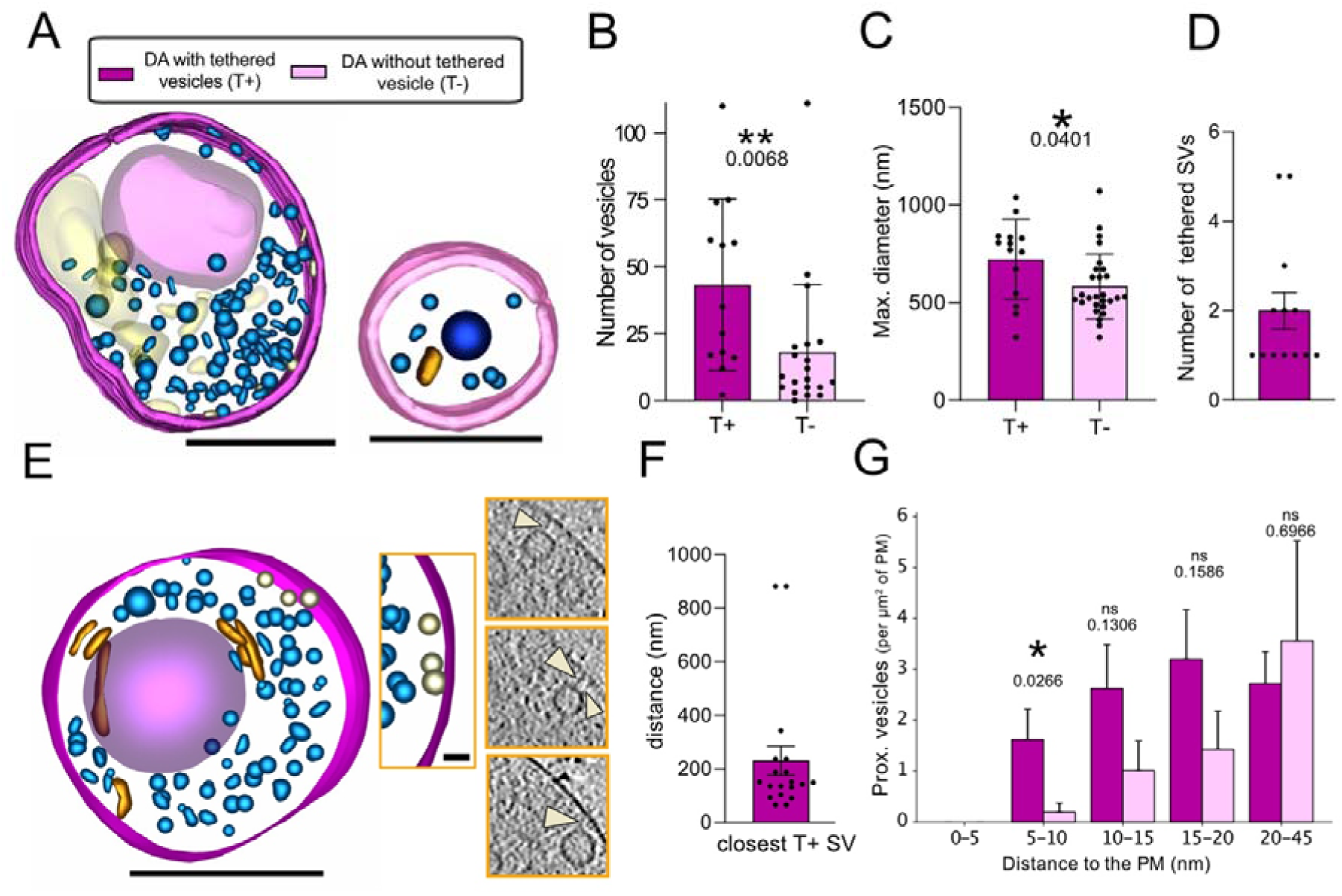
Distribution of tethered vesicles in DA synaptosomes. **A**, 3D models of DA synaptosomes with (T+, left) and without (T-, right) tethered vesicles. Scale bars, 500 nm. Color coding as in Figure 1. **B**, Total number of vesicles in DA synaptosomes with (T+) or without (T-) tethered SV (Mann-Whitney; p-value = 0.0068). **C**, Diameter of DA synaptosomes with (T+) or without (T-) tethered vesicles (Mann-Whitney; p-value = 0.0401). **D**, Number of tethered vesicles per T+ DA synaptosomes. **E**, 3D model of a DA synaptosome with a cluster of 3 tethered SVs on one side of the plasma membrane. Scale bar 500 nm. On the right, detail of the synaptosome with the 3 tethered vesicles and corresponding tomogram images. Scale bar 50 nm. **F**, Nearest neighbour distance between tethers on the plasma membrane. **G**, Number of vesicles per µm^2^ of plasma membrane at increasing distances from the plasma membrane for T+ and T-DA synaptosomes (t-tests; p-values = 0.0266; 0.131; 0.159; 0.697).

We also compared DA synaptosomes forming CS-DHS (that is adhering to a GLU terminal) or not. We found no significant differences in their vesicle number, spatial organization and tethering (Figure S9). This suggests that the adhesion to a GLU terminal does not affect the propensity of DA synaptosomes to contain tethered vesicles.

### Correlation of DHS connection with alterations of CS terminals

We wondered whether the adhesion of a DA terminal to a GLU synapse affects the ultrastructure of the GLU presynapse. We compared GLU synaptosomes that were not part of DHS (GLU DA-synapses, Figure 6A, n = 30) with GLU synaptosomes engaged into a DHS (GLU DA+ synapses or DHSs, Figure 6B, n = 32), both from the dual color model. The total number of vesicles are not different between GLU DA- and GLU DA+ terminals, on average 205.7 and 222.9 vesicles (Figure 6C; p-value = 0.864), as well as, the density of vesicles, 1893 per µm^3^ and 1943 per µm^3^, respectively (Figure 6D; p-value = 0.577). Similarly, the AZ areas are not different (Figure 6E). Together, these argue that the morphology of terminals and overall distribution of SVs is the same.

**Figure 6:**
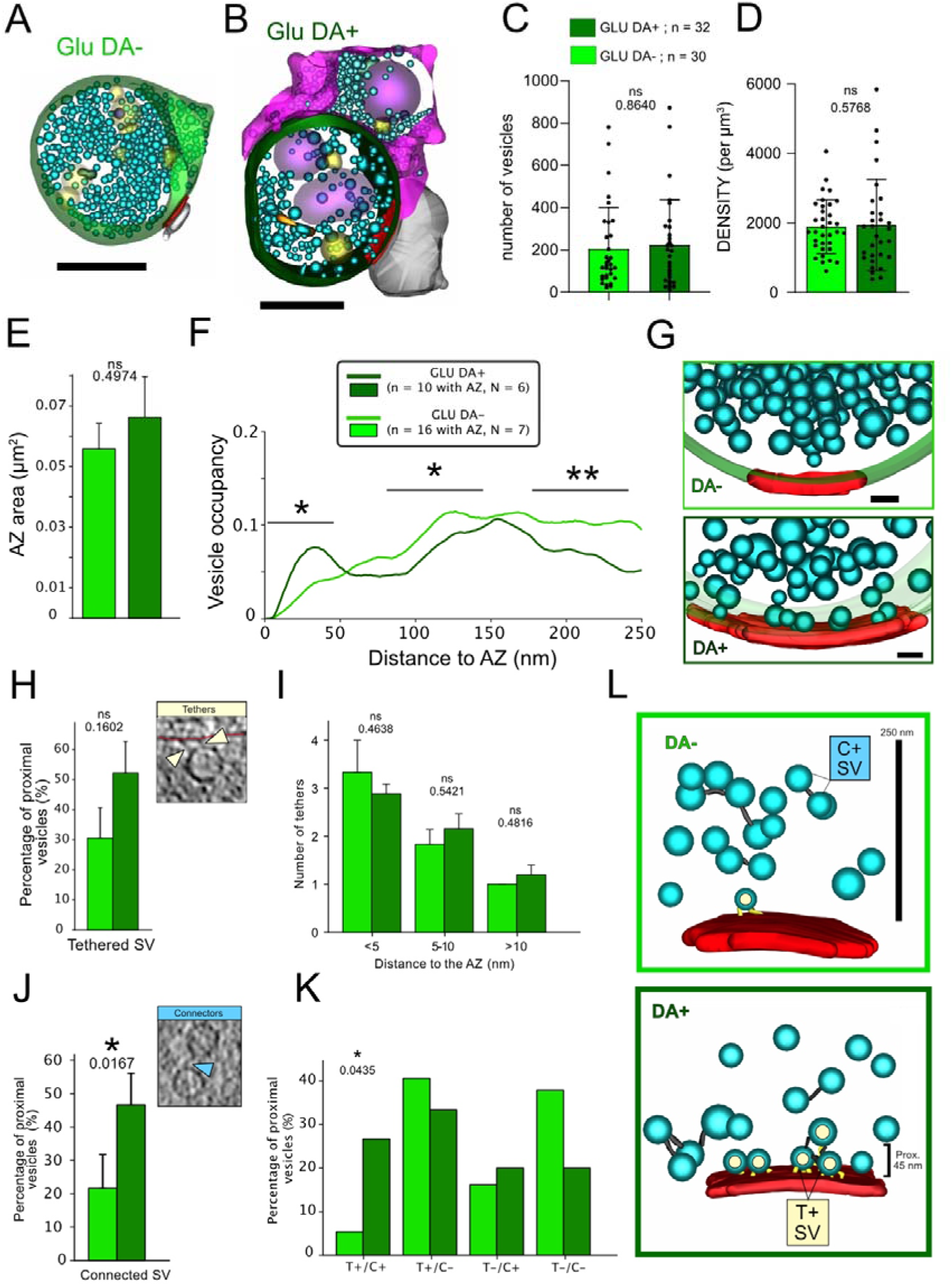
Comparison of GLU synaptosomes connected or not to a DA terminal. **A-B**, 3D models of a DA-GLU synaptosome (A) and a DA+ GLU synaptosome or DHS (B). Scale bars 500 nm. **C-D**, Number (C) (Mann-Whitney; p-value = 0.864) and density (D) of vesicles in DA- and DA+ GLU synaptosomes (Mann-Whitney; p-value = 0.577) **E**, Active zone area of DA- and DA+ GLU synaptosomes (t-test; p-value = 0.497). **F**, Average fraction of volume occupied by SVs vs distance to the plasma membrane for DA- and DA+ GLU synaptosomes (t-tests; p-values: 0 to 45 nm = 0.0215; 75 to 150 nm = 0.0195; 150 to 250 nm = 0.0016). **G**, 3D models of DA- and DA+ synaptosomes showing the number of proximal vesicles at the active zone. **H**, Percentage of proximal vesicles (< 45 nm from plasma membrane) that are tethered in DA- and DA+ GLU synaptosomes (t-test; p-value = 0.1602). **I**, Number of tethers per vesicle at various distances from the active zone (t-tests; p-values: <5 nm = 0.4638; 5 to 10 nm = 0.5421; >10 nm = 0.4816). **J**, Percentage of proximal vesicles that are connected to other vesicles in DA- and DA+ GLU synaptosomes (t-test; p-value = 0.0167). **K**, Proportion of proximal vesicles that are tethered and/or connected in DA- and DA+ GLU synaptosomes. The proportion of tethered and connected vesicles is significantly higher in DA+ GLU synaptosomes (chi-squared test; p-value = 0.0435). **L**, 3D models of DA- and DA+ GLU synaptosomes. In the DA-GLU synaptosome, a single SV is tethered and not connected to other vesicles. In the DA+ GLU synaptosome, four SVs are tethered and connected to one more vesicle.

Neverthless, the fraction of the volume occupied by proximal SVs was significantly higher in GLU DA+ than in GLU DA-synaptosomes, and showed a peak that signifies a higher concentration of proximal SVs, while the occupancy peak was absent in GLU DA-terminals (Figure 6F,G). A precise characterization of SV location and tether length showed that the results for GLU DA+ terminals were similar to those of unperturbed hippocampal glutamatergic synapses reported previously ^3^, while GLU DA-terminals had a lower proportion of SVs located 5-10 nm to the AZ membrane but, surprisingly, a higher proportion of tethers 6-12 nm in length (Figure S10).

Additionally, proximal vesicles in GLU DA+ synaptosomes are significantly more connected to other vesicles than in GLU DA-synaptosomes (46.7% vs. 21.6%; p-value = 0.0167) (Figure 6J). The fraction of tethered vesicles was about 70 % bigger in GLU DA+ terminals but did not reach significance. However, the fraction of proximal SVs that are both tethered and connected was significantly higher in GLU DA+ terminals (26.7% vs. 5.4%; p-value = 0.0435) (Figure 6K,L). Together with the observation that the number of tethers per SV was similar in GLU DA+ and GLU DA-terminals at all distances to the plasma membrane (Figure 6K), our results argue that connectors are primarily responsible for the observed difference in the proximal SV distribution.

## DISCUSSION

Here, we report the ultrastructure of fully hydrated, close-to-native DA terminals from adult mouse striatum at a single nanometer resolution, as well as of DHS, which comprise DA terminals in contact with GLU synapses and were characterized previously with immunofluorescence ^25^.

Synaptosomes obtained from mouse brain constitute a reliable model to investigate the spatial configuration of synapses ^16,38^. They are particularly amenable for cryo-ET because they can be directly observed by cryo-EM without further processing such as cryo-sectioning or cryo-focused ion beam milling ^16^. Moreover, they retain functionality, such as SV exocytosis and endocytosis, protein composition and post-synaptic calcium signalling ^3,29,35,38^. Likewise, we show with FM4-64 labelling that our preparation of synaptosomes undergoes stimulation dependent SV cycling. Moreover, the GLU synaptosomes we obtained from the mouse striatum are qualitatively and quantitatively comparable to forebrain and hippocampal synaptosomes (likely glutamatergic), and dissociated cultures observed previously with cryo-ET ^3–5,29,30^. The presynaptic element contains hundreds of small round SVs (∼40 nm diameter) and contact a PSE with a clearly defined PSD and a 32 nm wide synaptic cleft filled with dense material. This defines an AZ of ∼0.08 µm^2^, in the range of reported sizes varying between 0.04 µm^2^ and 0.10 µm^2^ ^3,48,49^. Moreover, SVs are polarized towards the AZ, with a peak of volume occupancy ∼25 nm from the plasma membrane, which reflects the enrichment in tethered and primed vesicles, similar to forebrain synapses and was proposed to be necessary for proper neurotransmitter release ^50^. Nevertheless, this peak was less pronounced in our sample of CS synaptosomes than in forebrain synaptosomes ^3,4^, which may reflect genuine differences in relative SV pool sizes in cortical/hippocampal vs striatal synapses. Remarkably, proximal SVs located less than 5 nm from the AZ have in both CS and cortical synapses on average 3 tethers ^4,5^, which likely corresponds to the primed state of SV and to functionally defined readily releasable SVs. In neurons lacking Munc13-1 and Munc13-2, which abolishes priming ^51^, proximal SVs have only 1 tether ^4^. Therefore, we propose that CS synapses have primed vesicles with the same structural hallmark as the ones defined in hippocampal synapses.

The most striking feature of DA synaptosomes is the relative sparsity of small synaptic-like vesicles, about 3-fold less dense than in GLU synaptosomes. Because DA synaptosomes are also smaller than GLU synaptosomes, there are 10 times less vesicles in DA synaptosomes than in GLU synaptosomes. These observations are consistent with ultrastructure of DA axons determined with serial electron microscopy reconstruction in chemically fixed striatum tissue in which axonal varicosities display very heterogenous ultrastructure: some have only few vesicles, others have small synaptic-like vesicles, larger ones, or both types of vesicles ^15^. Likewise, recent studies have shown qualitatively similar results in cultured DA neurons observed with cryo-CLEM ^52^ and DA neurons derived from human induced pluripotent stem cells ^53^. Vesicles in DA synaptosomes are also significantly more elongated than vesicles in GLU synaptosomes. Elongated-shaped vesicles have also been documented at GABAergic inhibitory synapses with conventional electron microscopy ^54^ and also with cryo-CLEM ^30^. Moreover, SVs in GLU synapses become elongated in the absence of VGLUT1, which reflects a lower luminal osmotic pressure ^55,56^. Therefore, DA vesicles, like GABA vesicles, experience different osmotic pressure than GLU vesicles.

Importantly, we show that, similar to GLU synaptosomes, vesicles in DA synaptosomes are frequently linked together by connectors and to the plasma membrane by tethers. Nevertheless, only 39 % of the DA synaptosomes contain at least one tethered vesicle. The DA synaptosomes with at least one tethered vesicle are bigger and contain twice as many vesicles as the ones without any tethered vesicle. Interestingly, other studies have shown that only 30 % of DA axonal varicosities contain active zone proteins bassoon, RIM or ELKS ^12^, and this proportion is also found for functional DA varicosities able to release fluorescent dopamine analog ^14^. Therefore, we propose that synaptosomes containing tethered vesicles correspond to release-competent DA terminals. Moreover, the number of tethers linked to proximal vesicles (less than 45 nm from the plasma membrane) is the same in GLU and DA synaptosomes. This suggests that similar molecules are involved in vesicle tethering in both types of synapses. Indeed, proteomic analysis of striatal DA synapses show similar degree of enrichment of the major proteins involved in vesicle tethering ^25,57^. However, DA synaptosomes lacked very proximal vesicles (less than 5 nm from the plasma membrane), which have on average 3 tethers, a hallmark of vesicles primed for release in GLU synapses. This difference points to a fundamental difference between these two types of terminals. We thus predict that the kinetics of vesicle exocytosis will be much slower in DA neurons than in GLU neurons, where SV exocytosis occurs in less than 1 ms after calcium entry. So far, the kinetics of DA vesicle exocytosis has been determined with fast amperometry or fluorescence imaging of dopamine sensors ^7,8^.The rise time of these events is greater than 10 ms but it could be dominated by diffusion of ligand to the detector micrometres away from the dopamine release site. More precise investigation of dopamine release is needed to determine if the kinetics of dopamine release is genuinely slower, and whether this is due to differences in vesicle priming.

We found a clear PSE at only 2 out of 94 DA terminals. These are among the synaptosomes which contain many small round vesicles, and are thus indistinguishable from GLU synaptosomes. They may correspond to terminals of DA/GLU neurons co-expressing the vesicular glutamate transporter VGLUT2 ^41,42,58^. In the adult mouse, these terminals are mostly concentrated in the shell of the nucleus accumbens. Interestingly, in these axons VGLUT2 is segregated from the vesicular dopamine transporter VMAT2, suggesting that glutamate and dopamine release sites are segregated ^58^. In the other 92 DA terminals reconstructed, no clear PSE was identified.

We identified previously that DA terminals can strongly interact with other synaptic elements in a so-called DHS ^25^. In the 32 reconstructed DHS, we found that 28 directly adhere to a presynaptic VGLUT1 element and 4 contacting the GLU PSE. On the other hand, DA terminals, identified in conventional transmission EM with immuno-labelling of tyrosine hydroxylase, are as likely to contact the pre- or post-synaptic side of CS synapses ^22^. The selection of DHSs by cryo-fluorescence with the presynaptic marker VGLUT1-Venus likely biased our sampling towards interaction with presynaptic markers. Nevertheless, we could characterize both pre- and post-synaptic DA/GLU adhesion sites in their native state with cryo-ET. We found that the adhesion sites had similar areas, around 0.05 µm^2^, as well as the cleft sizes, around 12 nm. However, these adhesions did not define a privileged tethering site for DA vesicles. Overall, we did not identify a localized site for DA vesicle tethering, suggesting a large AZ for exocytosis.

Finally, we showed that the morphology of GLU terminals and overall distribution of SVs are very similar in GLU DA+ and GLU DA-terminals. Also, we detected tethers of different lengths, which were previously associated with different molecular composition of tethers ^3^, in both types of presynaptic terminals. These suggest that the GLU terminal formation and their protein composition does not depend on the presence of DA terminals. Nevertheless, the distribution of proximal SVs in GLU DA+ terminals was consistent with those of non-perturbed glutamatergic synapses, while GLU DA-terminals showed a flat proximal SV distribution profile previously associated with a reduced neurotransmitter release ^3,4^. Furthermore, we correlated these differences with changes in SV connectivity and possibly also with tethering. Therefore, our data indicate that the contact between DA and GLU terminals at DHS modulates the SV organization at the single nanometer scale, as well as release properties of GLU terminals, and that this modulation is mediated by SV connectors. We could hypothesize that the formation of DHSs in the striatum is an important feature modulating glutamate release. Overall, DHSs could be an important substrate for the modulation of glutamate release and striatal activity in vivo. These multi-partite assemblies offer a close proximity between dopamine release sites and specific glutamatergic synapses. By this mechanism, the spatial and temporal synchronization between dopamine and glutamate activity would be maximal, which could play a major role in the plasticity of excitatory input to striatal neurons ^24^.

## Materials and Methods

### Animal models

Three mouse models have been used. The VGLUT1^venus^ knock-in line to label CS synapses. The DAT-cre BAC transgenic mouse line^33^ in which we transduced VTA/SNc neurons with an AAV1 carrying pCAG-Flex-mNeongreen coding sequences to label dopaminergic neurons projecting to the striatum (coordinates from bregma are A/P: 2.9mm; M/L: 1.6 mm; D/V: 4.6 mm with 12° angle). To co-detect CS synapses with DA terminals in the same sample, we used a double labelling approach. DAT-cre crossed with the reporter Ai14TdTomato mouse line were crossed with VGLUT1^venus^ KI mice. Adult mice of both genders, aged 12 to 18 weeks were used.

We refined the experimental design and the procedures to reduce as possible the number of animals used and their suffering. All procedures were in accordance with the European guide for the care and use of laboratory animals and approved by the ethics committee of Bordeaux University (CE50) and the French Ministry of Research under the APAFIS n° #21132 and #38144.

### Preparation of synaptosomes

The preparation of synaptosomes was adapted from a previously published protocol ^59^. Briefly, animals were euthanized by cervical dislocation, decapitated and the head was immersed in liquid nitrogen for 5 seconds for rapid cooling but not freezing of the tissue. The striata were subsequently dissected under an epi-fluorescence stereomicroscope (Leica Microsystems, Germany). Samples were then homogenized in 1.5 ml of ice-cold isosmolar buffer (0.32 M sucrose, 4 mM HEPES pH7.4, protease inhibitor cocktail Set 3 EDTA-free (EMD Millipore Corp.)) using a 2 ml-glass-Teflon® homogenizer with 12 strokes at 900 rpm. The homogenate (H) was centrifuged at 1000 g for 5 min at 4 °C in a benchtop microcentrifuge. The supernatant (S1) was separated from the pellet (P1) and centrifuged at 12500 g for 8 min at 4 °C. The crude synaptosomes pellet (P2) was suspended in 350 μL of isosmolar buffer and layered on a two-step ficoll density gradient (5 mL of 13% Ficoll and 5 mL of 7.5% ficoll, both in 0.32 M sucrose, 4 mM HEPES). The gradient was centrifuged at 50,000 × g for 1 h and 10 min at 4 °C (Optima L100XP Beckman Coulter with SW32Ti rotor). The synaptosome fraction was recovered at the 7.5 and 13% ficoll interface using a 0.5 ml syringe. An additional centrifuge of the collected fraction was performed in 2ml of HEPES-buffered Krebs like solution (HBK) ^36^ composed of : 143 mM NaCl, 4,7 mM KCl, 1,3mM MgSO_4_, 1,2 mM CaCl_2_, 20 mM HEPES, 0,1mM Na_2_HPO_4_ and 10 mM D-glucose, pH = 7,4) at 12,500g for 5min in order to wash the excess of ficoll and sucrose residues. The barely visible pellet was resuspended in 200µl HBK leftover and placed for 15 minutes at 37 °C for physiological recovery before plunge-freezing. Alternatively, S1 fraction diluted in HBK has been used allowing for faster preparation with comparable results.

### FM4-64 uptake and release assay

To assess synaptosome integrity, we performed a fluorogenic vesicle exo-endocytosis assay using the amphiphilic styryl dye FM4-64 (Thermo Scientific T13320). Synaptosomes were prepared fresh with the same protocol as for cryo-CLEM. Then they have been diluted in HBK and centrifuged 34 min at 6750g on 12 mm coverslips coated with 1 mg/ml Poly-L-Lysine. Imaging was performed right after on a wide field epifluorescence microscope Leica DMI8 equipped with an inverted 63X/1.4 oil objective, a Hamamatsu Flash 4.0 v2 camera and a controlled 37°C/CO2 chamber. Coverslips were placed in a recording chamber (Ludin), incubated with 500 µl of prewarmed HBK and imaged in TRITC and GFP channels at different registered positions over a 5 µm stack (1^st^ acq). HBK was removed and FM4-64 (6 ng/µl final concentration), KCl (40 mM final concentration) and 37°C/CO2 HBK mixture was added inducing exo-endocytosis cycle and staining membrane and internalized vesicles (see supplementary figure 1A). After 1 min 30 s, KCl was rinsed twice with HBK and FM4-64 with HBK was added staining the remaining plasma membrane. After 1 min 30 s, second acquisition was launched (2^nd^ acq). After 5 rinses with HBK, a third acquisition captured signal from internalized vesicles only (3^rd^ acq). KCl-HBK mixture was added to induce depolarization and exocytosis and distaining was imaged 1 min 30 s after (4^th^ acq). Loading was measured by subtracting FM intensity of single synaptosome at acquisition n°3 (vesicles stained) versus acquisition n°1 (background). Release was measured by subtracting intensity of single synaptosome at acquisition n°3 (vesicles stained) with acquisition n°4 (non exocytosed vesicles and background dye binding). Control experiments with no KCl for loading or no KCl for release were performed. Analysis was performed on Fiji^60^ using a home-built macro-command.

### Plunge-freezing

Quantifoil R2/2 Cu 200 or R3.5/1 Cu 200 mesh grids (Q-R2_2-2C100 or Q-3.5_1-3C100 Delta Microscopies) were glow-discharged for 30 seconds at 2.5 mA using a ELMO glow discharge system (Cordouan Technologies) to enhance their hydrophilicity. Following this, 4 µl of freshly purified synaptosomes, mixed with with 10 nm colloidal gold beads (Sigma 741957) and 100 nm FluoSpheres^TM^ beads (F8797 ThermoFischer) were applied to the grids. The excess sample was immediately blotted away from the opposite side for 5 seconds in a Vitrobot Mark IV (Thermo Fisher Scientific), maintained at a temperature of 4 °C and at 100 % humidity. Finally, grids were plunge-frozen in liquid ethane and stored in liquid nitrogen until observation.

### Cryo-fluorescence microscopy

We used a commercially available system to perform cryo-fluorescence microscopy (Leica – DM6 FS Cryo CLEM). We monitored on test grids with TetraSpecks^TM^ beads (Thermo Scientific T7279) shifts between channels and drift artifacts in cryogenic conditions. No significant shift was noticed, while noticeable mechanical drift occurs for acquisitions of more than 20 z-steps, which may be necessary when the grid is not flat and exactly parallel to the focal plane. Therefore, we limited our acquisitions to flat areas and imaged consecutive z planes with an increment of 0.5 to 1 µm (20 z-steps max). A custom-made Plexiglas® chamber was built around the set-up to maintain relative humidity under 40% reducing contaminations by water. Temperature of the objective chamber was maintained at -190°C (83K) using a pump projecting vapor of fresh liquid nitrogen. Each grid was observed separately reducing the time spent in the chamber to 25 minutes maximum. Sample was observed with a 50X objective (Leica – HC PL APO 50x/0.90NA Dry 11566064) and z-stacks were acquired through a Hamamatsu ORCA-Flash 4.0 camera, in 4 channels, in the following order: Brightfield (Empty); Tx Red (Em: BP560/40 nm; Ex: BP630/75 nm); GFP (Em: BP470/40 nm; Ex: BP525/50 nm); DAPI (Em: BP360/40 nm; Ex: BP470/40 nm). Large, flat area of the grids were acquired using the LAS X Navigator mosaic imaging tool. Grids were stored in liquid nitrogen until the cryo-EM session.

Images from regions of interest were stacked in a maximal intensity Z-projection using Fiji software for each fluorescence channels ^60^. Minimal Z projection was performed for the brightfield channel allowing the detection of holes in the carbon layer. Contrast has been adjusted and allowed detection of all synaptosomes (dim and bright), as well as fluorescent beads. TIFF images have been saved in PNG file format for editing. Each square of interest (with one or more synaptosome in a hole) was identified and marked with a unique number. Cropping of each corresponding area into a single image was done and resulting PNG files were saved-back into TIFF files for compatibility with Serial EM ^61^.

### Correlation between fluorescence and electron microscopy

The correlation process was carried out in 2 stages. The initial “approximate” correlation focused on locating the grid squares captured through cryo-fluorescence microscopy. After inserting the grid into the cryo-transmission electron microscope, a low-magnification montage (34x) was constructed using SerialEM ^61^ to visualize the entire grid. The fluorescence image was then imported into the software via the “Import Map” function. Areas of interest were identified using obvious landmarks, such as carbon holes and the center of the grid, which were visible in both imaging modalities.

Once the squares were identified, a high-magnification montage (1600x) was created for each area of interest. The corresponding fluorescence image for each square was imported separately. Precise correlation in SerialEM was done using the fluorescent fiducial beads as registration points, detectable in both fluorescence and electron microscopy. Beads were selected and assigned specific identifiers on the fluorescence image, and the same beads were subsequently located on the electron microscopy image. A total of 5 to 10 surrounding beads were used for the transformation to correlate with a single target for tomography. Both images were then correlated using the “Transform Item” function.

### Cryo-electron tomography

Tilt series of the synaptosomes were recorded using a Talos Arctica microscope (Thermo Fisher Scientific), operating at 200 kV and equipped with a K2 Summit direct electron detector (Gatan). The tilt series were acquired in a bidirectional scheme using SerialEM 4.0, covering angles from -60° to +60° with 2° increments, at a magnification of 11,000x, resulting in a pixel size of 3.987 Å. Imaging was performed with an underfocus of -8 µm, and the cumulative electron dose ranged between 80-90 e⁻/Å².

Tilt series alignment and tomogram reconstruction were carried out using EMAN2 ^62^ for quick visualization and data selection. IMOD software package 4.11.25 ^37^ integrated into the SCIPION framework ^63^ was used to reconstruct selected tomograms for segmentation in 3dmod (see below). Reconstruction was achieved using weighted back-projection combined with a SIRT-like filter. Binning of 4 was applied and resulted in a pixel size of 1.5588 nm.

### Segmentation and production of models

We selected tomograms with accurate correlation, sufficient contrast and intact synaptic features for segmentation (numbers provided in Table1). Other tomograms were discarded (examples in Figure S8).

Manual segmentation was performed using 3dmod, a software from the IMOD package ^64^. Synaptosome plasma membranes were segmented until disappearance and meshed, therefore no interpolation was used and missing wedge volume was not corrected for the synaptosomes. Organelles and vesicles were interpolated using spherical interpolation, resulting in closed objects. Active zone area was defined as plasma membrane portion facing the post-synaptic density. Contact area (DHS) was the plasma membrane portion of the DA synaptosomes following tight apposition (< 15 nm) to the GLU plasma membrane. Synaptic vesicles were defined as objects smaller than 80 nm with round or elongated shapes. Mitochondria were easily detected as dense folded membranes are present in the lumen. Vesicular bodies were observed thanks to the presence of one or more vesicle-like structure inside. Objects with folded, irregular shapes and bigger diameter have been categorized separately and correspond to unidentified structures.

### Data analysis

We obtained geometrical information such as volume and maximum diameter from the 3D models using *imodinfo* command lines (IMOD package). Sphericity was calculated using Wadell’s index ^65^ (W_i_) with the following formula 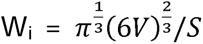 where V represents the volume and S the surface of the vesicle. Thus W_i_ = 1 for a perfect sphere and decreases for non-spherical objects.

Vesicle density inside synaptosomes was obtained dividing the number of vesicles by the volume available inside the synaptosome. In Fiji, cytoplasmic density profiles were obtained averaging a 40 nm stack and pixel intensities were measured using the built-in commands. In Fiji, synaptic cleft and interspace density profiles were measure at 3 different single z planes (low, middle and high) in the stack and averaged for each tomogram. Values were normalized to the neighboring non-cleft area corresponding to background. We excluded tomograms in which gold beads were present as it artificially alters the density.

SV tethers and connectors were detected in and automated, template-free manner using the hierarchical connectivity algorithm, and their morphology, localization and interrelationship was analyzed by Pyto package (version 1.10, available at https://github.com/vladanl/Pyto), as described before ^29,47^. Briefly, for the analysis of vesicle distribution (volume occupancy), the presynaptic cytoplasm (including SVs) was divided into 1-pixel-thick layers according to the distance to the AZ membrane, and the fraction of the layer volume occupied by SVs was measured. Connector and tether lengths were calculated as the minimal edge-to-edge distance between connector / tether voxels contacting an SV or plasma membrane that takes into account central regions of tethers and connectors. In this way, the ambiguity inherent to the measurement of length of 3D objects is resolved and curvature of tethers and connectors contributed to their calculated lengths. However, possible extended protein-lipid binding regions are not considered. All image processing and statistical analysis software procedures were written in Python and implemented in Pyto package [37]. Pyto uses NumPy and SciPy packages and graphs are plotted using Matplotlib^66–68^ Statistical analysis was performed between the experimental groups using only planned, orthogonal comparisons. For the analysis of properties pertaining to individual SVs, connectors and tethers (such as the SV distance to the AZ membrane, tether length and fraction of tethers/connectors having a certain property), values within experimental groups were combined. Bars on the graphs show mean values and error bars the standard error of the mean (sem). In cases a fraction of SVs or tethers is shown, the error bars represent sem between synapse means. We used Student’s t test for statistical analysis of values that appeared to be normally distributed (e.g., vesicle diameter) and K-W test (nonparametric) for values deviating from the normal distribution (e.g., number of tethers and connectors per vesicle). For frequency data (e.g., fraction of connected and non-connected vesicles), χ2 test was used. In all cases, confidence levels were calculated using two-tailed tests. The confidence values were indicated in the graphs by a single asterisk for P<0.05, double for P<0.01, and triple for P<0.001.

We focused on tomograms originating from the VGLUT1^venus^ x DAT-cre x Ai14 tdTomato because it is the only model where we could reliably identify both GLU and DA and thus determine CS-DHS and non-CS-DHS conditions with certainty (see Table 2). Moreover, we selected tomograms where the contrast was optimal and sufficient information was available. We picked GLU synaptosomes where we identified the active zone and for DA where there were more than 2 vesicles. Thus, it ensures an optimal detection and a reliable comparison of the filaments between both conditions. We normalized tomogram density and applied a Kernel Gaussian filter (sigma = 2) to improve detection. Tomograms used for the glutamatergic active zone were cropped on the dedicated region and analysis was performed blind to the DHS, non-DHS condition.

### Estimation for shortest distance between tethered vesicles

We measured the arc distance between two tethered DA vesicles d (that is the distance while remaining in the plasma membrane) by measuring the distance in the projection in the tomogram plane measured along the arc of the plasma membrane (d_xy_) and the distance between the two tomogram planes in which the tethers connect the plasma membrane (d_z_) as 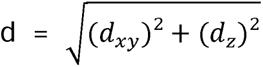. We estimated the distance between random points by taking the average maximum diameter of T+ DA synaptosomes D (722 ± 205 nm, Figure 5C). For the distance between random points, we approximated the projection of synaptosomes by a circle of radius R = D/2 = 361 nm. The average arc distance between random points in a circle is half the length of a half circle (by symmetry), or 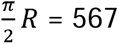 nm and its variance is 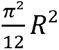 that is a standard deviation of 327 nm. This estimate is a lower limit of the true value, because it neglects the axial distance d_z_ and approximates synaptosomes as circles, minimizing distance.

## Acknowledgements

We thank Peter Vanhoutte and Nicolas Heck for providing the DAT-cre * Ai14 tdTomato line, the Pôle in Vivo for animal breeding, husbandry, help with stereotaxic injections. We thank members of the Bordeaux Imaging Center, namely Monica Fernandez-Monreal from for help with cryo-fluorescence microscopy and Fabrice Cordelières for the analysis of FM4-64 loading experiments. We thank student interns Timothé Lapha, Jade Giraud and Solène Hospital for reconstruction of some of the tomograms.

This work was supported by ANR (DopamineHub ANR-19-CE16-0003 to E.H. and D.P., FrontoFAT ANR-20-CE14-0020-03 to E.H. and UltraDopa ANR-24-CE16-5973-01 to E.H., D.P. and R.F.), the Fondation Recherche Médicale (to P.L., end of PhD and D.P., FRM team), the European Research Council (ERC consolidator Grant PneumoTransfo to R.F.) and the Regional Council of Nouvelle Aquitaine (ParkSynGraft 205024) to E.H. and D.P. This work was supported by the French Government through the France 2030 program [grant number 21-ESRE-0024] managed by the French National Research Agency (ANR) as part of the "Investissements d’avenir" program.

## Author contributions

P.L. performed stereotaxic AAV injections. synaptosome preparations and live imaging of FM4-64 uptake with the help of V. P-B. P.L. performed synaptosome freezing, cryo-fluo imaging, cryo-EM and cryo-ET with the help of R.A. E.M. and R.F. supervised the cryo-ET methodology. P.L. did all reconstructions and annotations of tomograms under the supervision of E.H. and D.P. P.L. performed quantitative analysis of all data together with V.L. P.L., R.F., E.H. and D.P. acquired funding. P.L., E.H. and D.P. wrote the manuscript and all other authors edited it.

