## Supplemental Figures 1-10 Tables 1-2 for "Cryo-correlative light and electron tomography of dopaminergic axonal varicosities reveals non-synaptic modulation of cortico-striatal synapses"

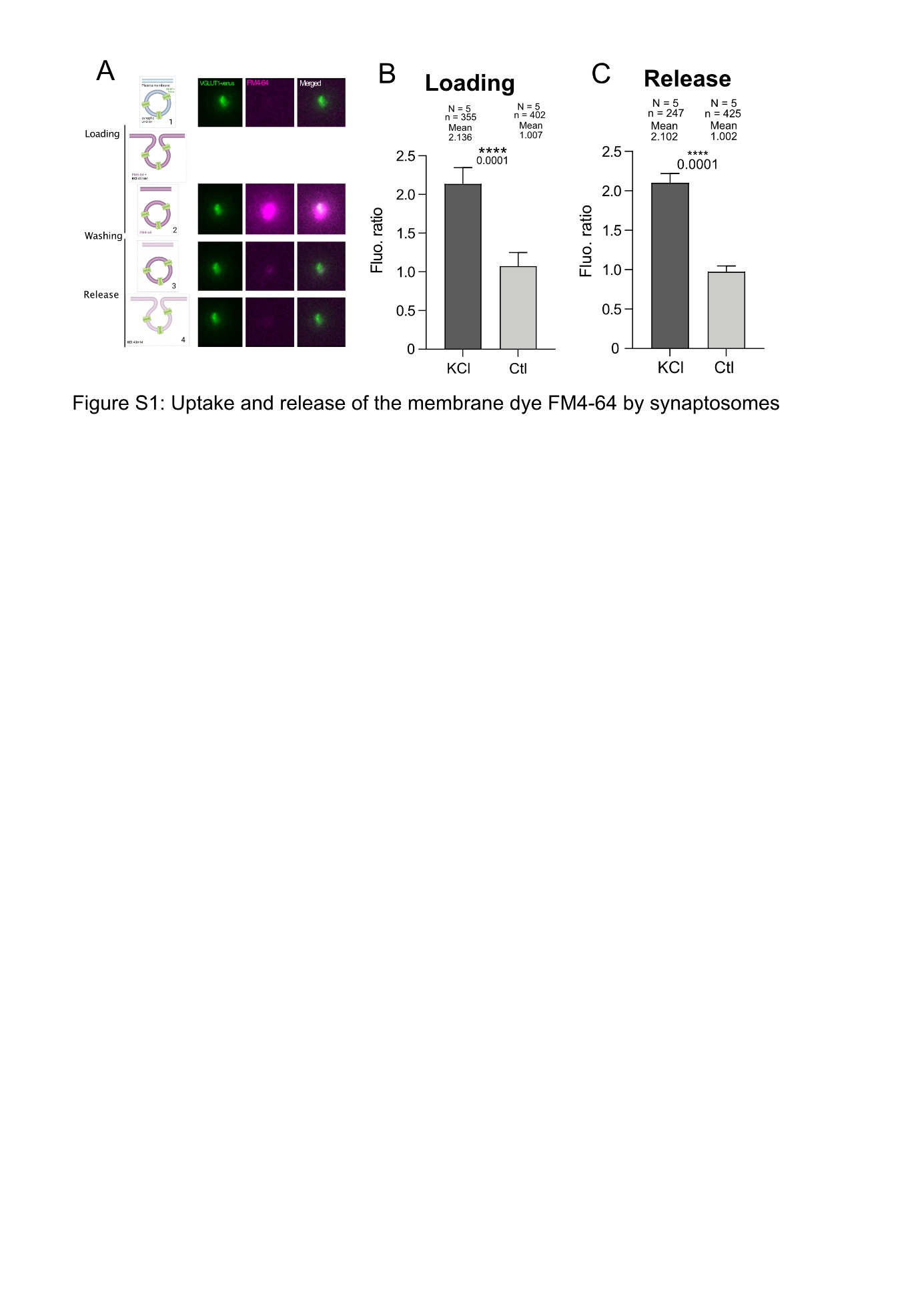

**Figure S1: Synaptic vesicle cycling in synaptosomes measured with FM4-64 uptake and release. A**, Scheme of the principle of FM assay in synaptosomes. A first image is acquired before adding the dye and corresponds to the background (1). FM4-64 is loaded into vesicles by triggering a first exo-endocytic cycle with 40 mM KCl and a second image is taken (2). After washing with HBK, only cycling vesicles remain stained by FM4-64 monitored by a third image (3). A second stimulation with 40 mM KCl induce release of the dye and is followed by washing with HBK and acquisition of a final image (4). A control experiment is performed without KCl depolarization for loading to monitor unspecific dye labelling. **B**, Violin plot showing the distribution of loading values (acq. 3 intensity - acq. 1 intensity) normalized to the control. On average the loading signal is 2.136 times higher upon stimulation with KCl (dark grey) which correspond to synaptosomes that have loaded the dye (Mann-Whitney; p-value < 0.0001). **C**, Violin plot showing the distribution of release values (acq. 3 – acq. 4) after stimulation (dark grey) normalized to the control (light grey). Destaining averages 2.102 times higher than the mean value of control (Mann-Whitney; p-value < 0.0001).

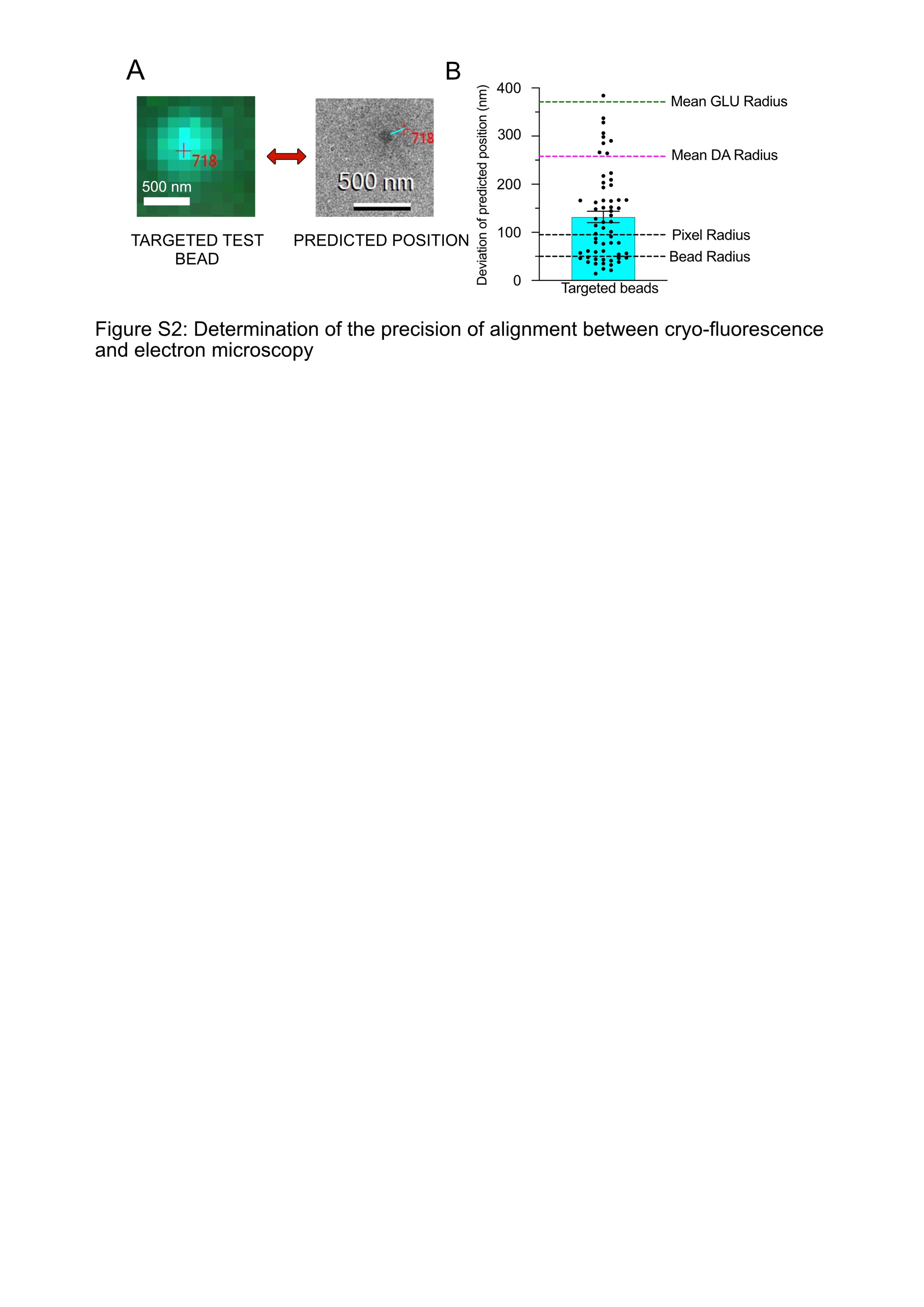
**Figure S2: Pointing precision of the alignment between fluorescence and electron microscopy A**, Representative images of a fluorescent fiducial bead that has not served for the correlation, in cryo-fluorescence microscopy (left) and in cryo-EM (right). The red cross (718) indicates the targeted position on the fluorescent image and the predicted position obtained on the cryo-EM after transformation. The cyan line shows the distance between the predicted position and the actual bead **B**, Histogram showing the deviation of the predicted positions in nanometers for beads that have not been used for the transformation. The average distance is 116 nm which corresponds to 44% of the mean DA radius and 31% the mean GLU radius (n = 64).

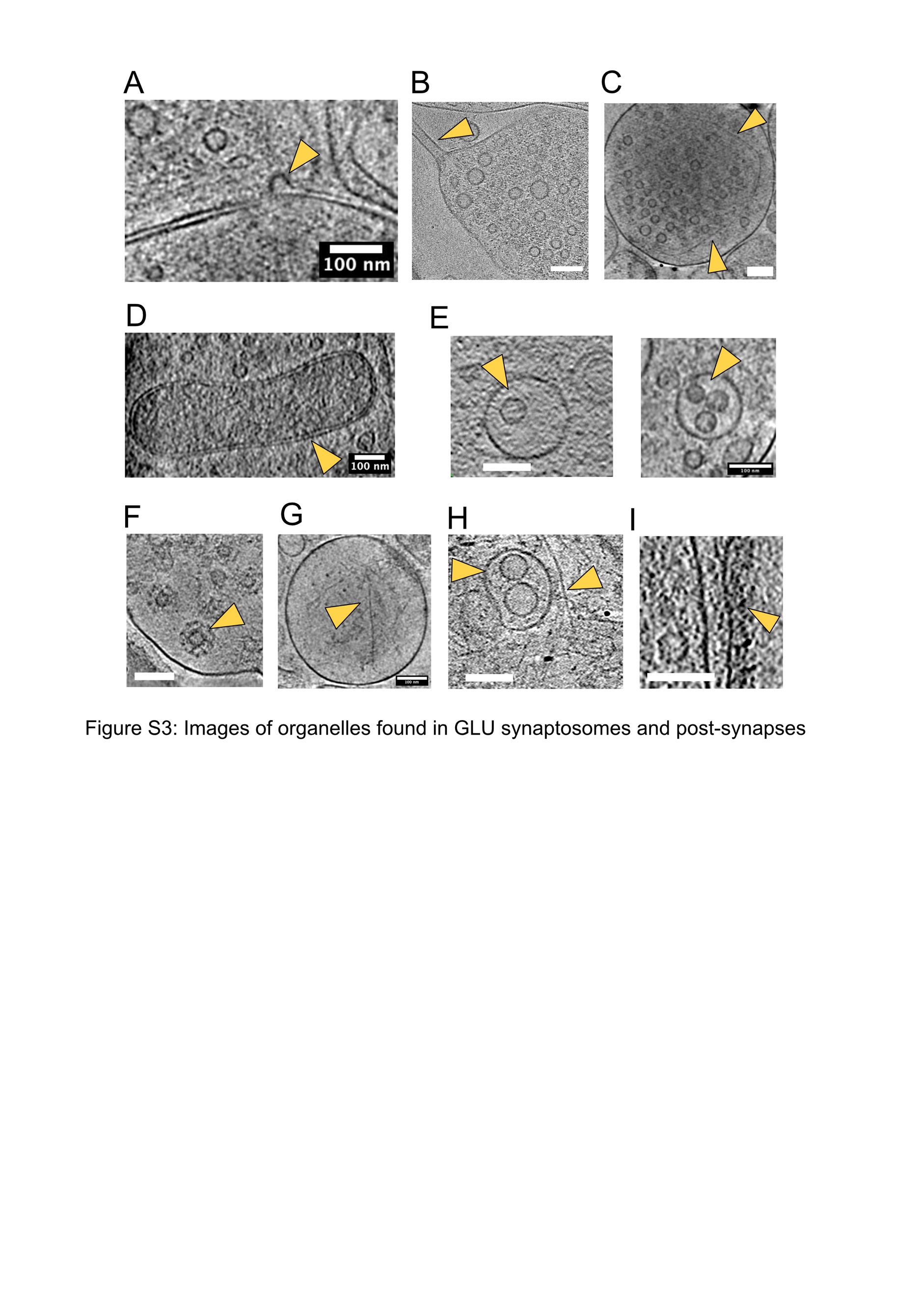

**Figure S3: Examples of structures and organelles found in GLU synaptosomes (A-F) and post-synaptic elements (G-I).** Orange arrows point to the organelles of interest in all panels.  **A**, Example of a membrane invagination, either an exocytic or endocytic event. **B**, Example of a narrow membrane tubule on one side of the GLU synaptosome. Perhaps a remaining axonal part filled with microtubules. **C**, Example of filaments surrounding synaptic vesicles. **D**, Example of a mitochondrion. **E**, Examples of multivesicular bodies. **F**, Example of clathrin-coated vesicles. **G**, Example of actin filaments in the PSE. **H**, Examples of a multivesicular body and a dense actin filaments network. **I**, Example of a post-synaptic density. Scale bars 100 nm.

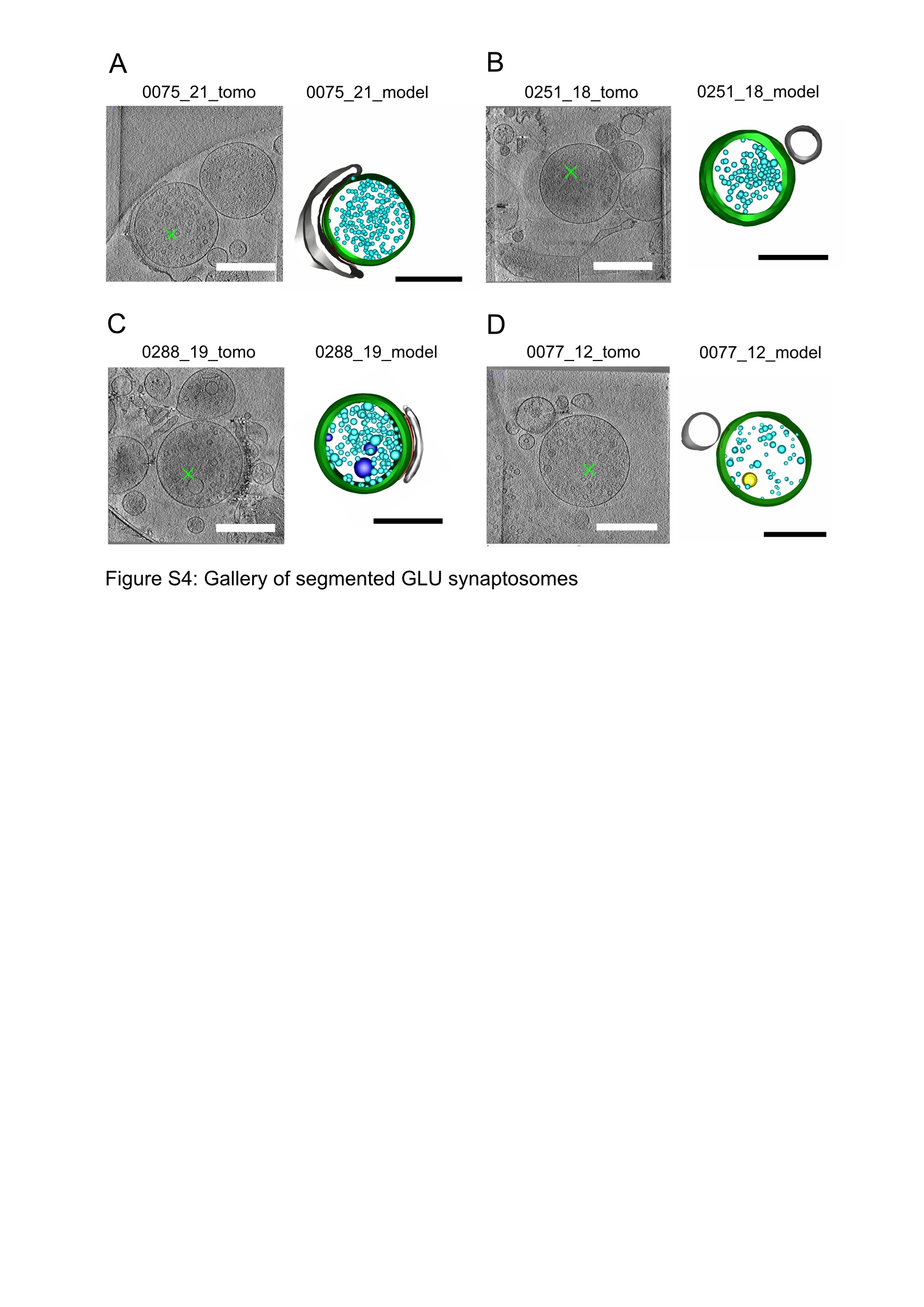

**Figure S4: Gallery of segmented GLU synaptosomes. A,** Example of a tomogram (numbered 0075_21) showing a GLU synaptosome (green cross, left) and the corresponding 3D model (right). The plasma membrane is represented in green, the post-synaptic element in light grey and synaptic vesicles in cyan **B,** Similar example for the tomogram numbered 0251_18. **C**, Similar example for tomogram 0288_19. Multivesicular bodies are drawn in dark blue. **D,** Similar example for tomogram 0077_12. Large vesicle is shown in yellow. Scale bar: 500 nm.

For the 3D models, the color coding is the same as in Figure 2 and 3.

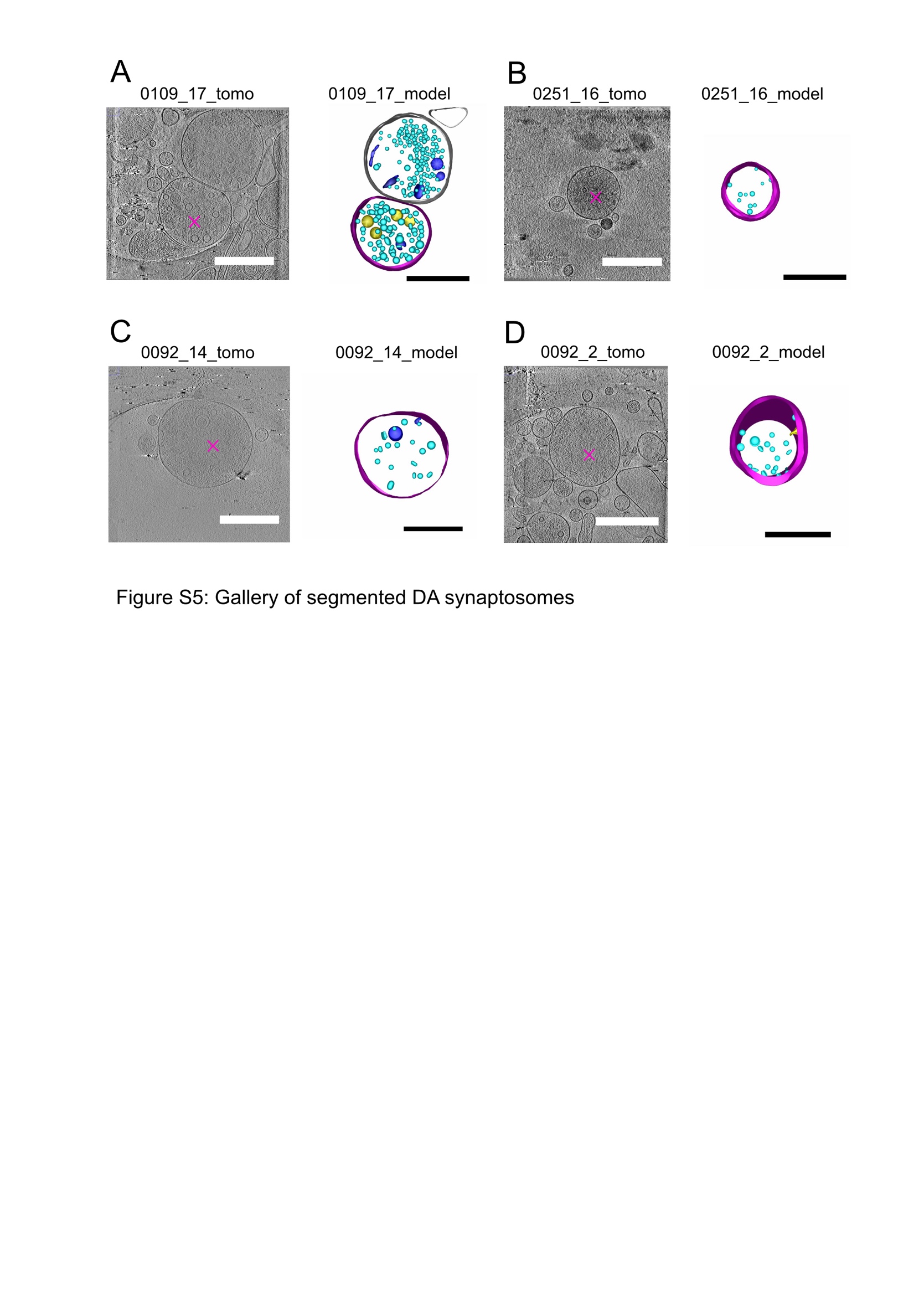

**Figure S5: Gallery of segmented DA synaptosomes. A,** Example of a tomogram (numbered 0109_17) showing a DA synaptosome (magenta star, left) and the corresponding 3D model (right). The DA element adheres to another element resembling a GLU synapse with many SVs and a clear PSE (top right). However, this tomogram was obtained in a mouse in which only DA neurons are fluorescent (DAT-Cre injected with AAV-Flex-NeonGreen), so the identity of the adhering structure could not be confirmed with fluorescence. **B,** Similar example for tomogram 0251_16 **C**, Similar example for 0092_14 **D,** Similar example for 0092_2. Scale bars: 500 nm. For the 3D models, the color coding is the same as in Figure 2 and 3.

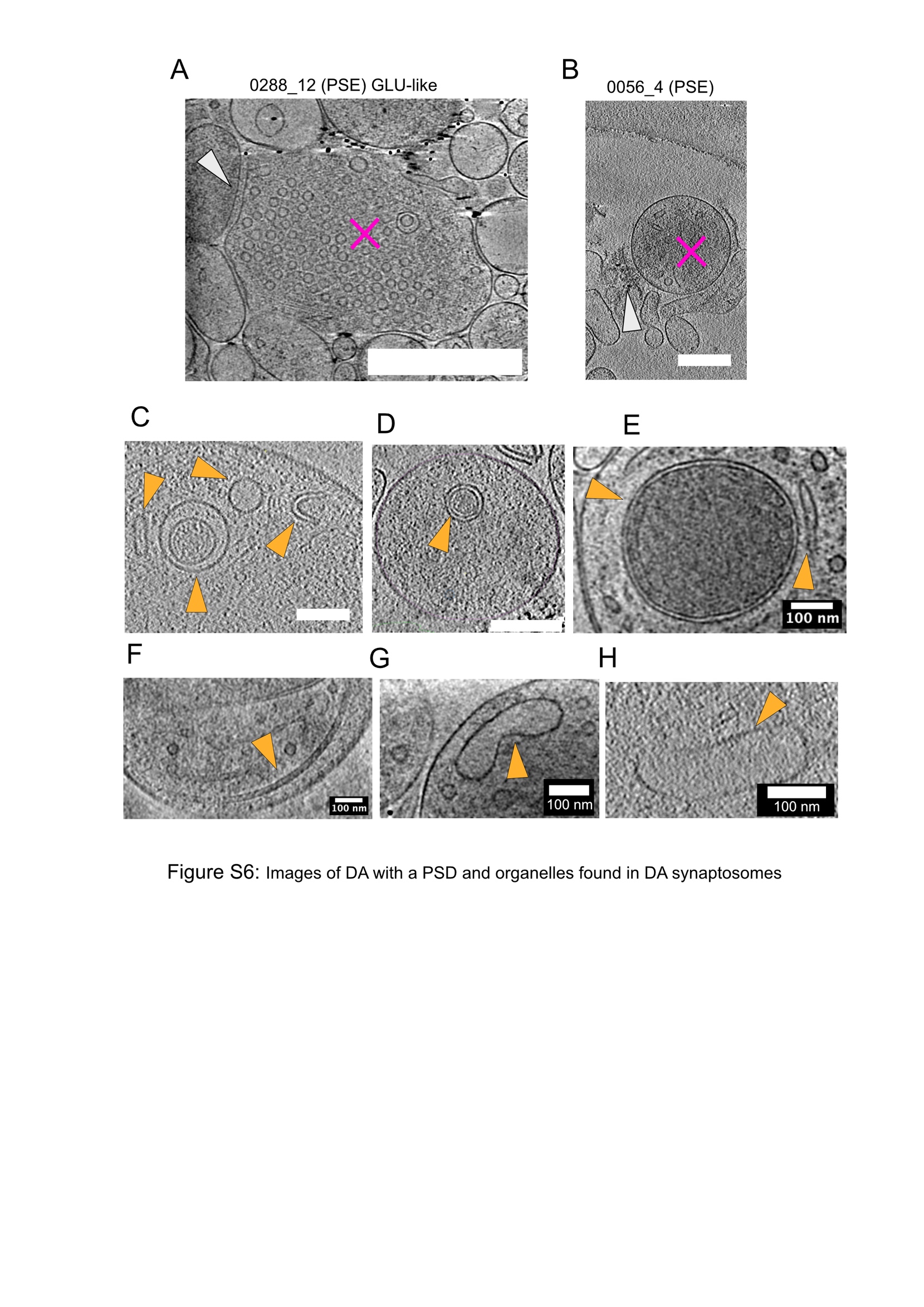

**Figure S6: Images of DA with a PSD and organelles found in DA synaptosomes. A**, DA synaptosome (magenta star) resembling a GLU synaptosome facing a PSE containing a PSD (grey arrow). It has 610 small synaptic vesicles, the highest observed in DA synaptosomes. Scale Bar: 500 nm. **B**, DA synaptosome (magenta star) facing an opened PSE with a PSD (grey arrow). Scale bar: 200 nm. **C**, Cytoplasmic content of a DA synaptosome containing, from left to right, a tubular vesicle, a vesicular body, a large vesicle, a C-shaped structure. Scale bar: 200 nm. **D**, A vesicular body. Scale bar: 200 nm. **E**, A mitochondrion (left) and an ER-like structure (right). Scale bar: 100 nm. **F**, A microtubule, Scale bar: 100 nm. **G,H**, Endosome-like structures. Scale bars: 100 nm.

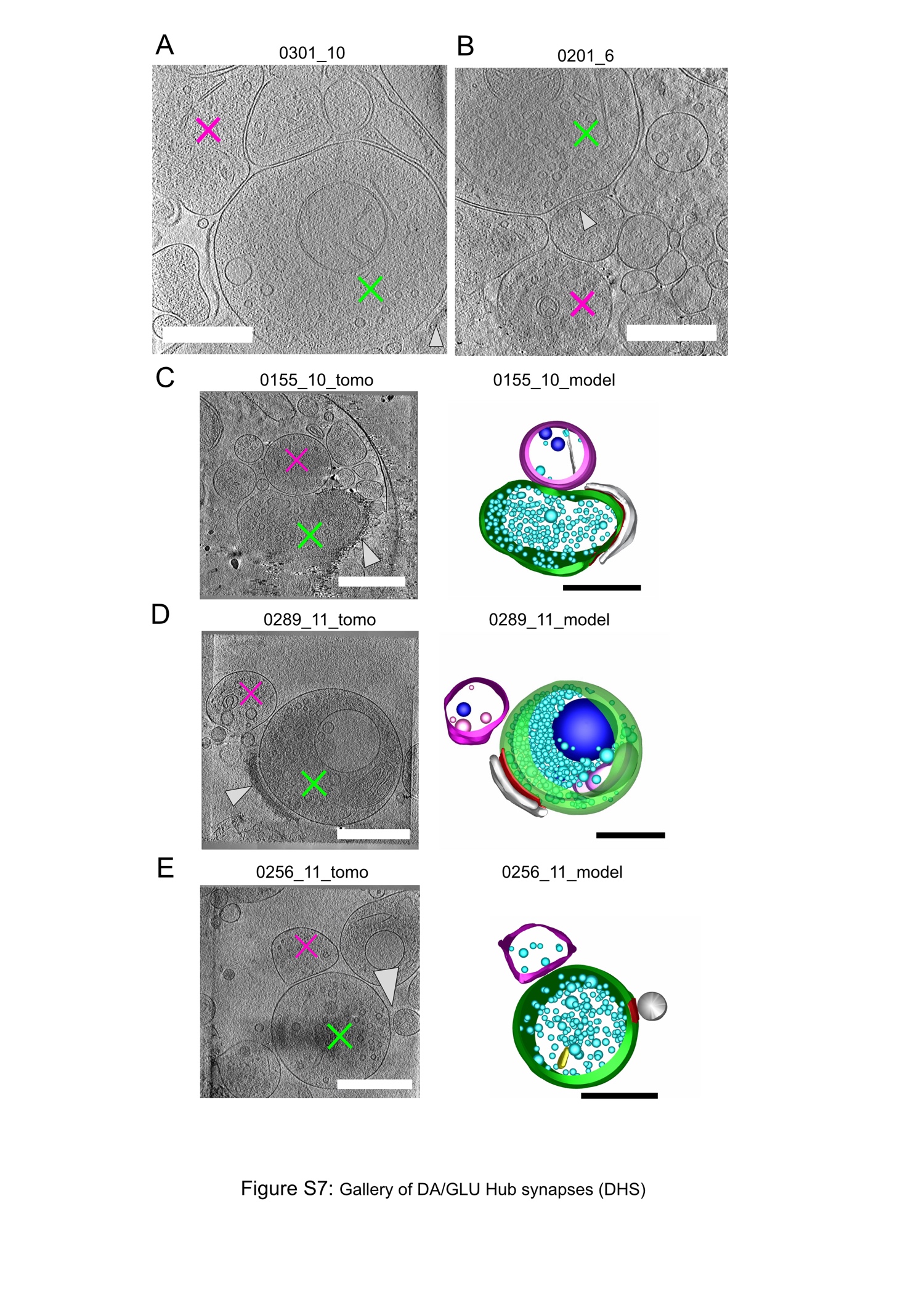

**Figure S7: Gallery of reconstructed CS-DHS identified with both DA and GLU reporters. A**, Single plane of the tomogram 0301_10 showing the DHS corresponding to the model in Figure 3B. DA is shown by the magenta star, GLU by the green cross and the post-synapse by the grey arrow. **B,** Single plane of the tomogram 0201_6 showing the DHS corresponding to the model in figure 3C. **C, D, E**, Examples of tomograms showing DHS (left) with their corresponding 3D model (right). Scale bars: 500 nm.

Color code is similar to fig S4,5.

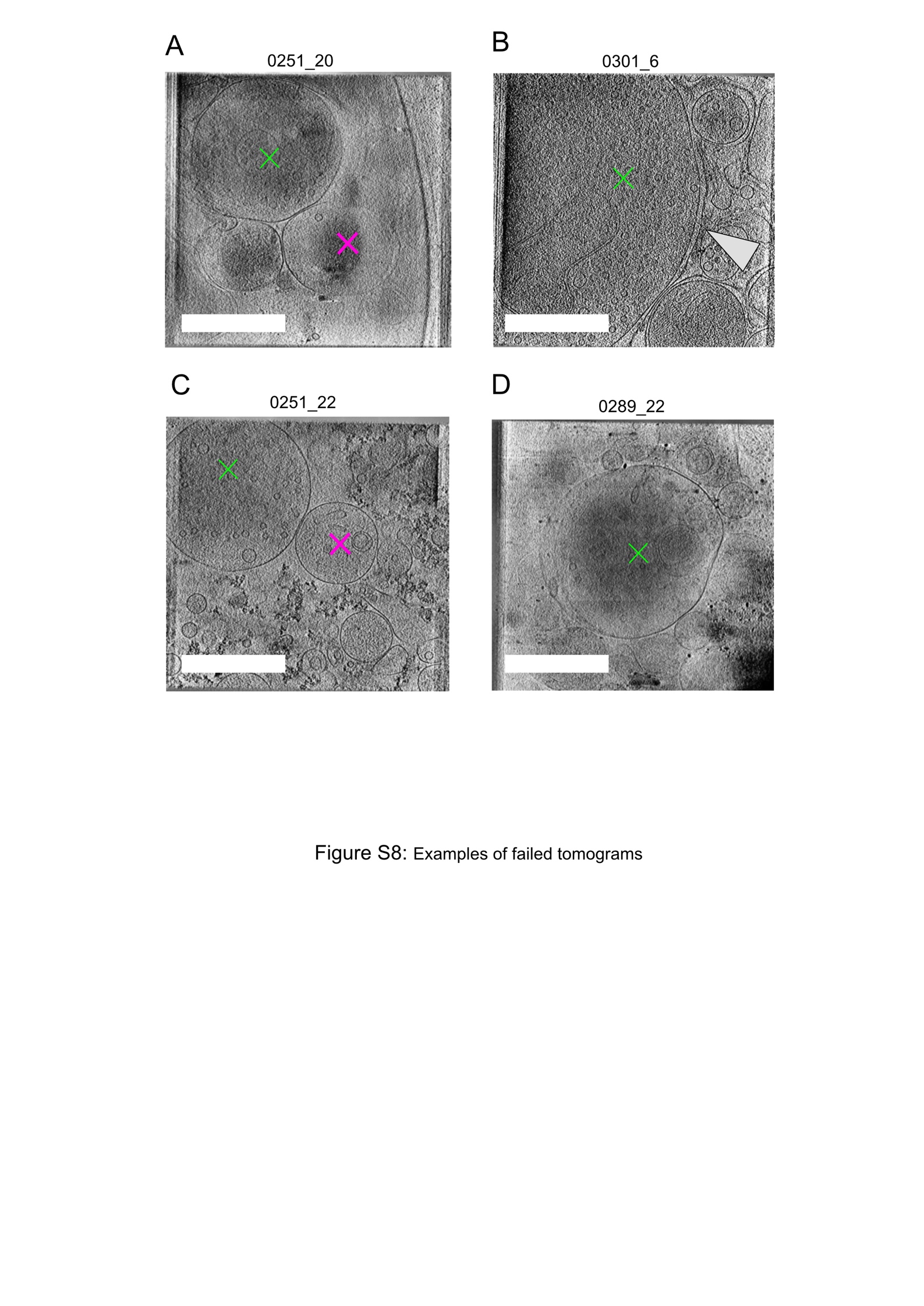

**Figure S8: Examples of tomograms not retained for segmentation. A**, Example of a tomogram where reconstruction resulted in a miss-alignment of the planes. **B**, Example of a tomogram plane showing a GLU synaptosome which is too big to obtain a sufficient contrast. **C**, Example of a tomogram where both DA and GLU structures are unrelated according to the correlation. **D**, Example of a tomogram for which the reconstruction failed, it results in very low information across the planes. Scale bars: 500 nm.

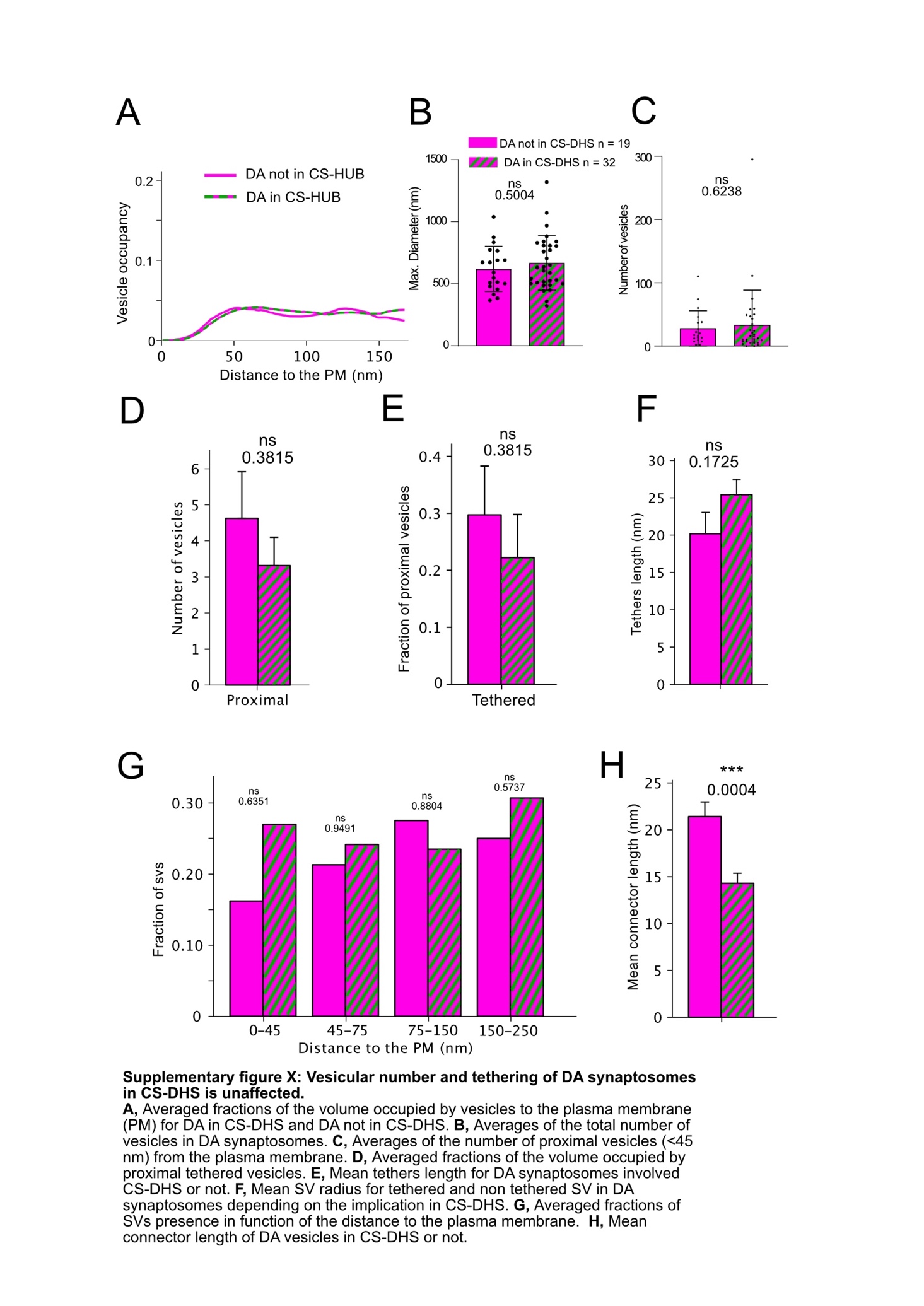

**Supplementary Figure 9: DA synaptosomes organization is unaffected in CS-DHS.** **A,** Averaged fractions of the volume occupied by vesicles to the plasma membrane (PM) for DA forming CS-DHS or not. **B,** Mean maximal diameter of DA synaptosomes involved or not in CS-DHS (Mann-Whitney; p-value = 0.500). **C,** Mean number of vesicles in DA synaptosomes forming CS-DHS or not (Mann-Whitney; p-value 0.624). **D,** Averages of the number of proximal vesicles (<45 nm) from the plasma membrane (Mann-Whitney; p-value = 0.382). **E,** Averaged fractions of the volume occupied by proximal tethered vesicles (t-test; p-value 0.382). **F,** Mean tether length in DA synaptosomes involved in CS-DHS or not (t-test; p-value = 0.173). **G,** Fractions of vesicles in function of the distance to the plasma membrane (t-tests; p-values: 0 to 45 nm = 0.635; 45 to 75 nm = 0.949; 75 to 150 nm = 0.880; 150 to 250 nm = 0.574). **H,** Mean connector length of DA vesicles involved in CS-DHS or not (t-test; p-value = 0.0004).

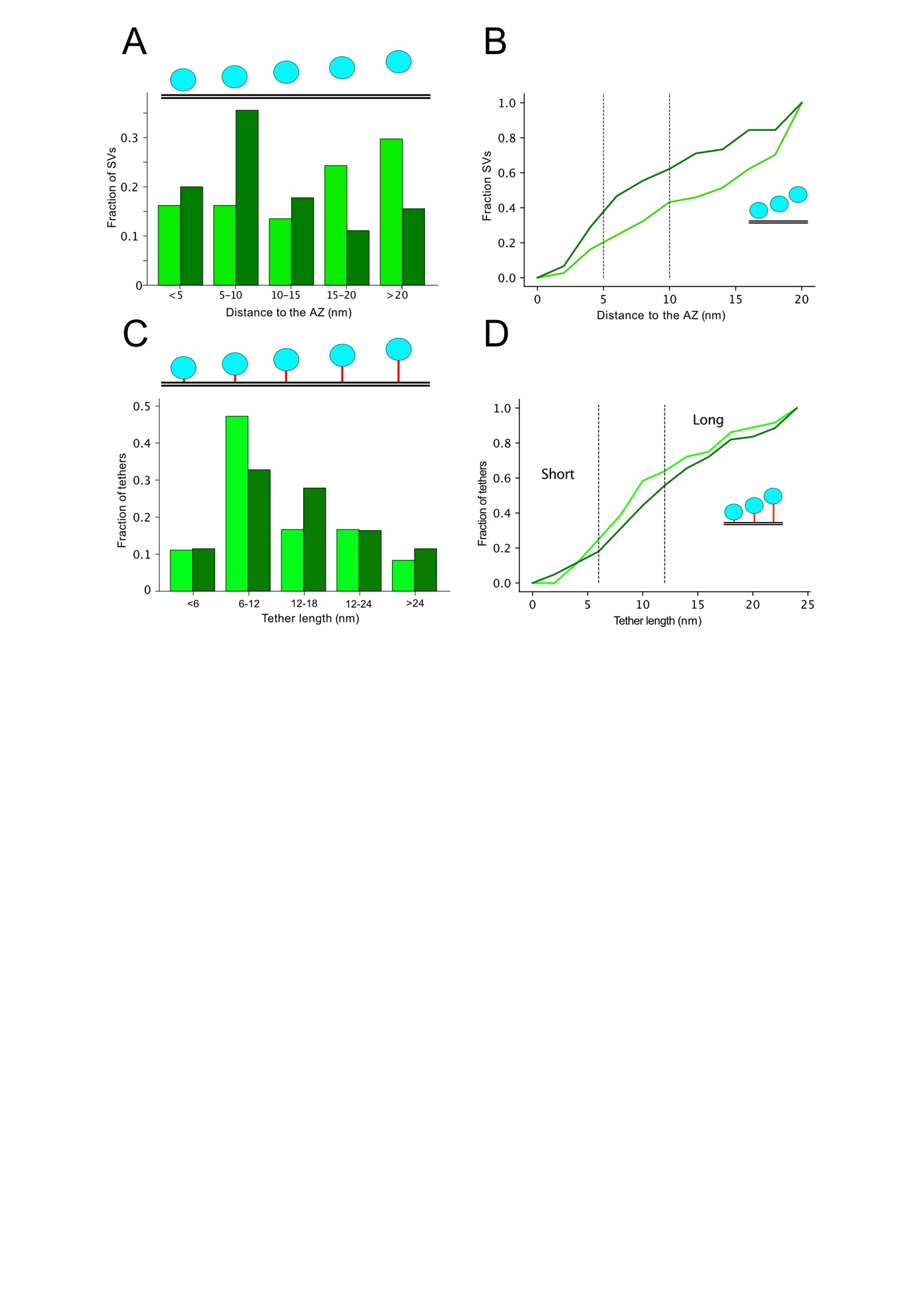

**Supplementary Figure 10: Distribution of proximal GLU vesicles in CS-DHS and tether length.**

**A,** Fractions of proximal vesicles (<45 nm) localized at certain distances from the active zone shown with histograms depending on the involvement in DHS (dark green) or not (light green). **B,** Cumulative distribution of the SV distances from the active zone, data are the same as in (A). **C,** Fractions of tethers corresponding to certain lengths. Tethers from GLU involved in DHS are in dark green, the ones not involved in DHS are in light green. **D,** Cumulative distribution of the length of the tethers. Data are the same as in (C).

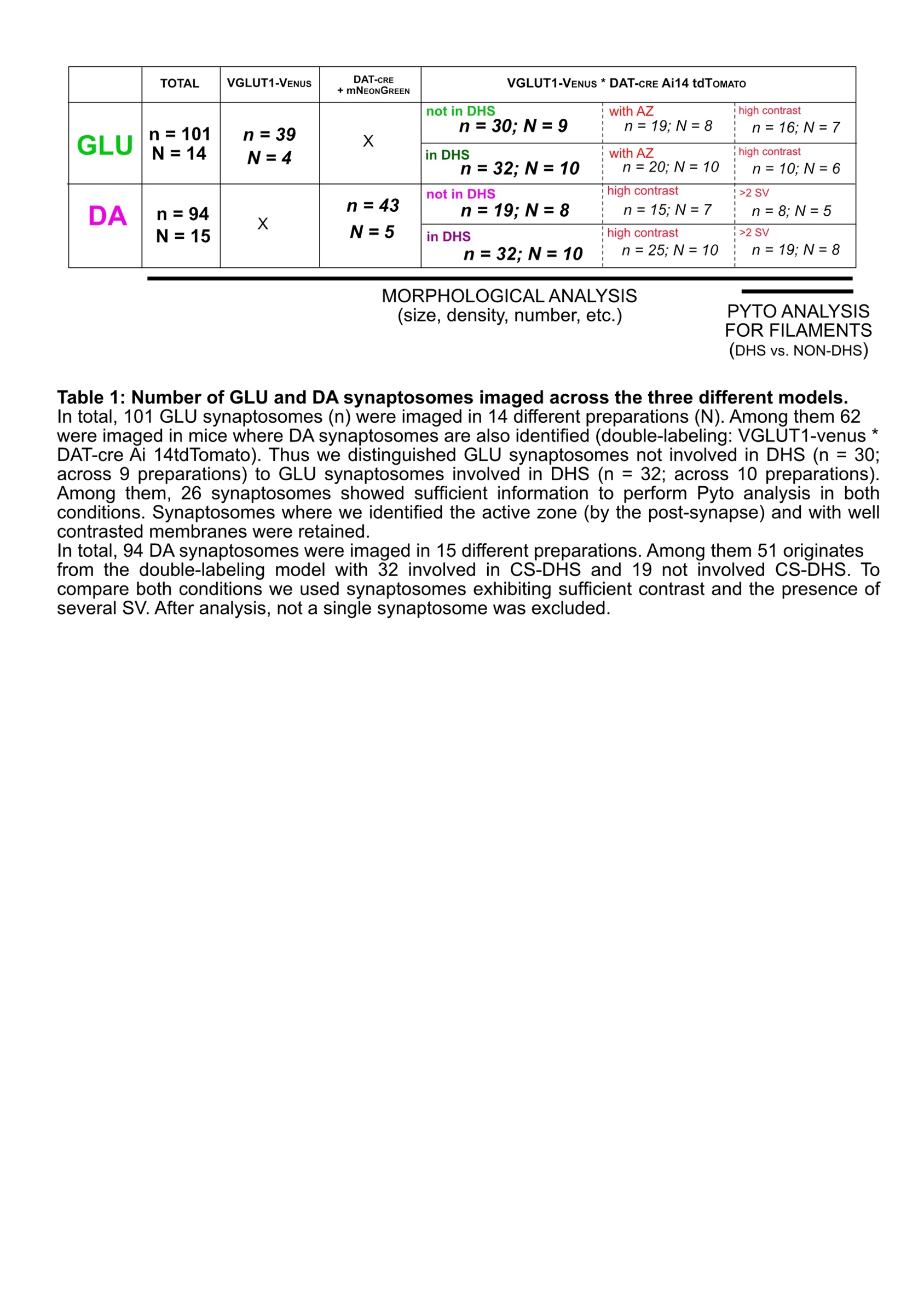

**Table 1: Number of GLU and DA synaptosomes imaged across the three different models.**

In total, 101 GLU synaptosomes (n) were imaged in 14 different preparations (N). Among them 62 were imaged in mice where DA synaptosomes are also identified (double-labeling: VGLUT1-venus * DAT-cre Ai 14tdTomato). Thus, we distinguished GLU synaptosomes not involved in DHS (n = 30; across 9 preparations) to GLU synaptosomes involved in DHS (n = 32; across 10 preparations). Among them, 26 synaptosomes showed sufficient information to perform Pyto analysis in both conditions. Synaptosomes where we identified the active zone (by the post-synapse) and with well contrasted membranes were retained.

In total, 94 DA synaptosomes were imaged in 15 different preparations. Among them 51 originates from the double-labeling model with 32 involved in CS-DHS and 19 not involved CS-DHS. To compare both conditions we used synaptosomes exhibiting sufficient contrast and the presence of several SV. After analysis, not a single synaptosome was excluded.

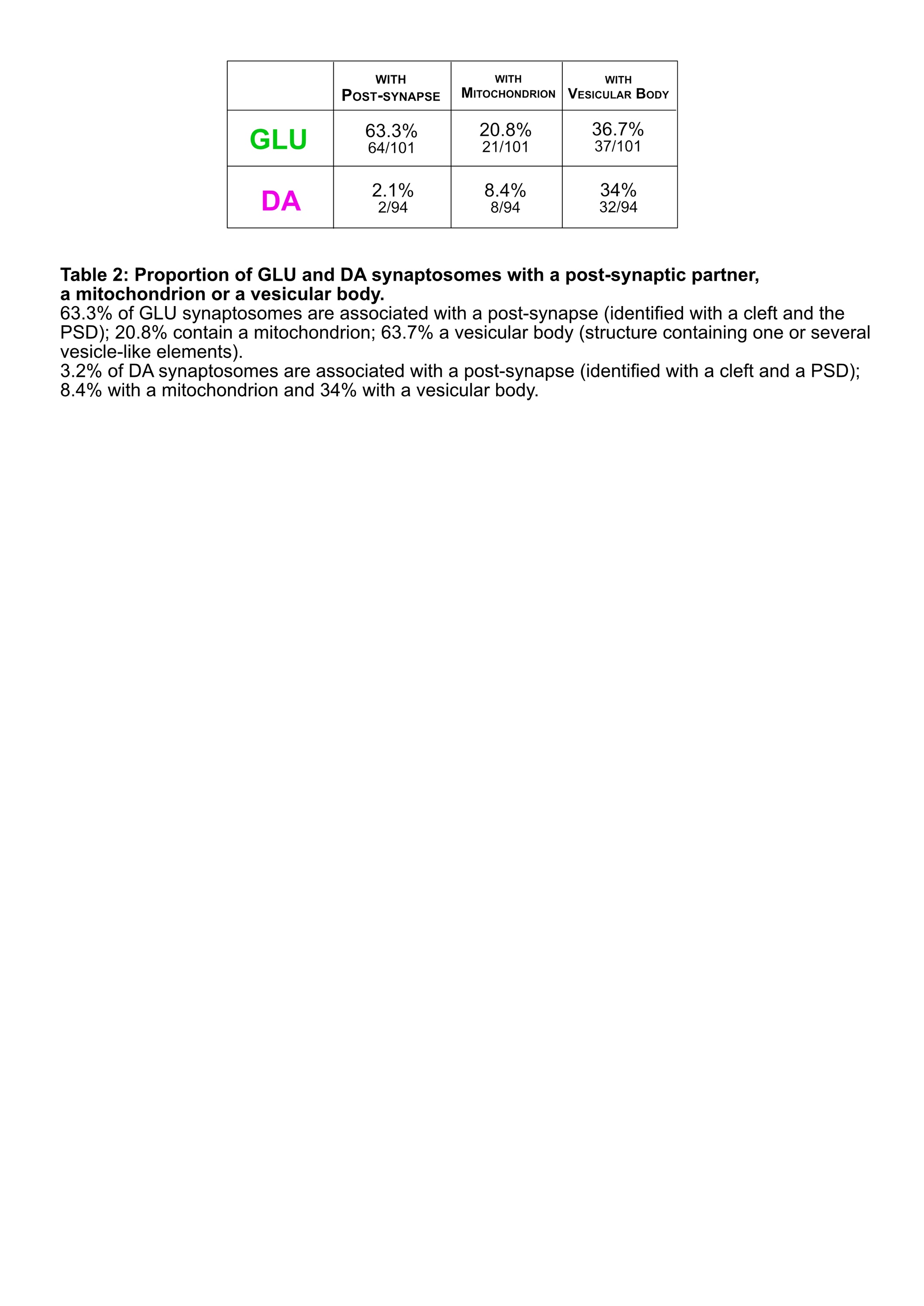

**Table 2: Proportion of GLU and DA synaptosomes with a post-synaptic element (PSE),**

**a mitochondrion or a vesicular body.**

63.3% of GLU synaptosomes are associated with a post-synapse (identified with a cleft and the PSD); 20.8% contain a mitochondrion; 63.7% a vesicular body (structure containing one or several vesicle-like elements). 2.1% of DA synaptosomes are associated with a post-synapse (identified with a cleft and a PSD); 8.4% with a mitochondrion and 34% with a vesicular body.
